# Radial glial lineage progression and differential intermediate progenitor amplification underlie striatal compartments and circuit organization

**DOI:** 10.1101/244327

**Authors:** Sean M. Kelly, Ricardo Raudales, Miao He, Jannifer Lee, Yongsoo Kim, Leif G. Gibb, Priscilla Wu, Katie Matho, Pavel Osten, Ann M. Graybiel, Z. Josh Huang

## Abstract

The circuitry of the striatum is characterized by two organizational plans: the division into striosome and matrix compartments, thought to mediate evaluation and action, and the direct and indirect pathways, thought to promote or suppress behavior. The developmental origins of and relationships between these organizations are unknown, leaving a conceptual gap in understanding the cortico-basal ganglia system. Through genetic fate mapping, we demonstrate that striosome-matrix compartmentalization arises from a lineage program embedded in lateral ganglionic eminence radial glial progenitors mediating neurogenesis through two distinct types of intermediate progenitors (IPs). The early phase of this program produces striosomal spiny projection neurons (SPNs) through fate-restricted apical IPs (aIP^*S*^s) with limited capacity; the late phase produces matrix SPNs through fate-restricted basal IPs (bIP^*M*^s) with expanded capacity. Remarkably, direct and indirect pathway SPNs arise within both aIP^*S*^ and bIP^*M*^ pools, suggesting that striosome-matrix architecture is the fundamental organizational plan of basal ganglia circuitry organization.

## INTRODUCTION

The striatum, the gateway and largest nucleus of the basal ganglia, receives massive inputs from all areas of the cerebral cortex and gives rise to the largest output circuits of the basal ganglia by way of multi-synaptic pathways leading to the brainstem and thalamus. Neural activity in the striatum itself is influenced by dopamine-containing afferents from the substantia nigra, serotonergic afferents from raphe nuclei, cholinergic inputs from the brainstem, and other neuromodulatory circuits. The striatum thus constitutes a key neural substrate through which diverse functionally specialized regions of the neocortex converge to modulate not only downstream motor programs for the initiation of voluntary behaviors, but also neural systems underpinning emotion, motivation, evaluation and learning (Graybiel, 2008; Grillner and Robertson, 2015; Hikosaka et al., 2014).

The majority of neurons in the striatum are output spiny projection neurons (SPNs). These are known to follow two fundamental organizational schemes. First, the striatum has a striking compartmental organization by which labyrinthine neurochemically specialized zones called striosomes (‘striatal bodies’) lie embedded within the much larger matrix compartment (Graybiel and Ragsdale, 1978). This striosome-matrix architecture contributes to the functional input and output connectivity of the striatum (Crittenden and Graybiel, 2011; Grillner and Robertson, 2015). The SPNs of striosomes receive preferential input from particular limbic regions and have privileged output to the dopamine-containing neurons of the substantia nigra and indirectly to the lateral habenula, thus engaging crucial dopamine- and serotonin-related neuromodulatory systems (Fujiyama et al., 2011; Jiménez-Castellanos and Graybiel, 1989; Watabe-Uchida et al., 2012; Crittenden et al., 2016; Eblen and Graybiel, 1995; Brimblecombe and Cragg, 2017). By contrast, SPNs of the much larger matrix compartment receive inputs from sensorimotor and associative cortical regions and in turn project to the main pallidonigral output nuclei of the basal ganglia that target downstream premotor regions and thalamocortical circuits (Donoghue and Herkenham, 1986; Flaherty and Graybiel, 1994; Graybiel, 2008; Hikosaka et al., 2014). Functionally, striosomal circuits have been implicated in evaluative functions related to affective control, cost-benefit decision-making, and reinforcement-based behaviors. By contrast, the matrix appears to be involved in the translation of cortical action plans and strategies to basal ganglia circuits involved in action execution (Amemori et al., 2011; Cui et al., 2014; Friedman et al., 2015; Fujiyama et al., 2011; Gerfen, 1992; Lopez-Huerta et al., 2015; White and Hiroi, 1998; Parthasarathy et al., 1992; Giménez-Amaya and Graybiel, 1991; Graybiel et al., 1994). The developmental basis of this striosome and matrix segregation is not well understood.

Superimposed upon striosome-matrix compartmentalization is a split between the projections of SPNs to the main basal ganglia output nuclei, the pallidum and the substantial nigra pars reticulata (SNr) (Gerfen, 1992; Kim and Hikosaka, 2015). SPNs projecting to the internal pallidum and SNr constitute the so-called direct pathway SPNs (dSPNs), conventionally considered to release basal ganglia-mediated inhibition or inhibition-excitation sequences and to promote action. By contrast, the SPNs projecting to the external pallidum give rise to the indirect pathway (iSPNs), considered to inhibit actions, potentially those with unwanted characteristics (Freeze et al., 2013; Albin et al., 1995). The direct-indirect pathway dichotomy has been elegantly formulated as a center-surround system in which wanted movements are promoted by the direct pathway and competing, interfering motor programs are inhibited (Mink, 1996)). By virtue of its large size and output organization, the matrix compartment is the main source of the direct and indirect pathways, but striosomes as well as matrix contain both dSPNs and iSPNs, at least as categorized by the expression of dopamine D1 (in dSPNs) and D2 (in iSPNs) receptors (Crittenden and Graybiel, 2011; Fujiyama et al., 2011; Brimblecombe and Cragg, 2017; Salinas et al., 2016).

Despite strong evidence for the coexistence of the striosome and matrix compartments and the direct and indirect pathways, remarkably little is known about the relationship between these two fundamental axes of striatal organization. A clue to the potential importance of developmental mechanisms underlying the compartmental divisions first came from studies demonstrating that SPNs making up striosomes and matrix have different birthdates, with striosomal SPNs being generated earlier than those within the matrix (Newman et al., 2015; van der Kooy and Fishell, 1987). Here we sought to link this evidence to neural progenitors and lineage mechanisms known to be crucial in striatal developmental cascades (Rubenstein and Campbell, 2013). The neuroepithelium of the lateral ganglionic eminence (LGE) of the embryonic ventral telencephalon consists of diverse types of progenitors that emerge and differentiate across different stages of striatal neurogenesis (Turrero García and Harwell, 2017). These include regenerative radial glial progenitors (RGs), which give rise to *apical* intermediate progenitors (aIPs) such as short neural progenitors (SNPs) and subapical progenitors (SAPs), and *basal* intermediate progenitors (bIPs) (Pilz et al., 2013). Although certain lineage relationships and neurogenic capacity of some of these progenitor types have been characterized by live imaging at mid-gestation (e.g., Pilz et al., 2013), their developmental trajectory through the course of LGE neurogenesis and especially their link to different SPN types and striatal circuit organization remain unknown. Tapping into the molecular mechanisms and transcription factors expressed in different progenitor types, we performed systematic genetic fate mapping to resolve different progenitor types and trace their developmental trajectories from early lineage progression to the distinct SPN types that they give rise to.

Here, we demonstrate that striosomal and matrix SPNs are sequentially generated from a RG lineage program through sequential allocation of distinct types of IPs with different amplification capacities. Whereas the early phase of this program produces striosomal SPNs through fate-restricted *Ascl1^+^* aIP^*S*^s with limited neurogenic capacity, the late phase amplifies matrix SPN production through the additional deployment and amplification of fate-restricted *Ascl1^+^*/*Dlx1^+^* bIP^*M*^s with expanded neurogenic capacity. During the final phase, *Dlx1/2^+^* bIPs specifically generate projection neurons of the annular compartment adjoining striosomes (A cells). Remarkably, a similar temporal and progenitor type split of striosomal and matrix SPN production was not observed for the genesis of D1- and D2-receptor bearing direct and indirect pathway SPNs. We found that each of these pathways is derived from both aIP^*S*^s and bIP^*M*^s throughout the course of RG lineage progression. These findings suggest that a primary lineage program within the LGE gives rise to striosome-matrix compartmentalization, and superimposed upon this organization, a secondary and different mechanism gives rise to direct and indirect pathway SPNs within each compartment. Thus the major components of striatal architecture are rooted in a radial glia lineage program differentiating future striosome and matrix compartments at the inception of striatal development, and they originate from sequential phases of lineage progression and from distinct intermediate progenitor types. These findings establish a novel framework for exploring the assembly of cortico-basal ganglia circuitry and should provide new clues to the differential vulnerabilities of the striosome-matrix and direct-indirect pathways in human disease states.

## RESULTS

### Early LGE Radial Glial Cells Sequentially Give Rise to Both Apical and Basal Intermediate Progenitors and to Both Striosomal and Matrix SPNs

The initial formation of LGE around embryonic day (E) 9.5 along the subpallium is followed shortly by the onset of neurogenesis, when proliferative neuroepithelial cells (NEs) begin to transform into neurogenic radial glial cells (RGs) (Sousa and Fishell, 2010). Although all striatal SPNs are thought ultimately to be generated from LGE RGs, recent studies indicate the existence of multiple types of progenitors with distinct molecular and morphological characteristics that are derived from RGs at different embryonic times (Pilz et al., 2013; Turrero García and Harwell, 2017). These include aIPs such as SNPs and SAPs), and also bIPs. How these different progenitor types relate to distinct striatal SPN types remains largely unknown. We used several inducible CreER drivers to carry out comprehensive genetic fate mapping of multiple LGE progenitor types.

Among the markers expressed in telencephalic progenitors (i.e., pallium and subpallium), the anti-proliferative protein TIS21 is unique in its specific expression in all progenitors undergoing neurogenic divisions, including subsets of RGs, aIPs, and bIPs (Attardo et al., 2008). To fate map different types of neurogenic progenitors, especially self-renewing RGs in the process of neurogenesis, we generated a knock-in *Tis21-CreER* driver and demonstrated that TM-induced activation of the *Ai14* reporter reliably and efficiently captured neuron-producing progenitors (Figures 1 and S1). The *Tis21-CreER* driver thus provided a unique and powerful fate-mapping tool for LGE neurogenic RGs starting from the very onset of neurogenesis. This enabled tracing of the RG lineage progression through sets of embryonic IPs to their SPN progenies in the postnatal striatum (Figures 1A and 1B). At the early onset of neurogenesis ~E10, 8-hour pulse-chase of Tis21^+^ progenitors labeled both RGs and aIPs (Figures 1C, 1F and 1G). We could not strictly distinguish between SNPs and SAPs, as these aIP subtypes are defined by their cell division patterns best captured by live imaging experiments (Pilz et al., 2013). Importantly, 48 hours after E10 TM induction (i.e., at E12), RFP^+^ RGs and aIPs remained abundant in the ventricular zone (VZ), indicating that RGs self-renewed and continued to generate aIPs. Sparse labeling allowed identification of elongated cell clusters, which likely represented clonal descendants from an initial neurogenic RG (Figure 1D, H, I; Figure S1D). Remarkably, even 96 hours after E10 TM induction (i.e., at E14), RFP^+^ RGs still remained in VZ, indicating persistent self-renewal (Figures 1E, 1J-1L, S1E, and S1F). Particularly large cell clones were frequent, which consisted of a “founder RG”, multiple Ascl1^+^ IPs, and larger number of postmitotic neurons likely engaged in migration along the RG radial fiber (Figures 1L and S1F, Supplemental Movie 1). Whereas E10-labeled RGs gave rise to aIPs from E10-E12 (Figures 1D, 1H and 1I), they made a transition to generating mainly bIPs in the subventricular zone (SVZ) by E14 (Figures 1E and 1J-1L).

**Figure 1:**
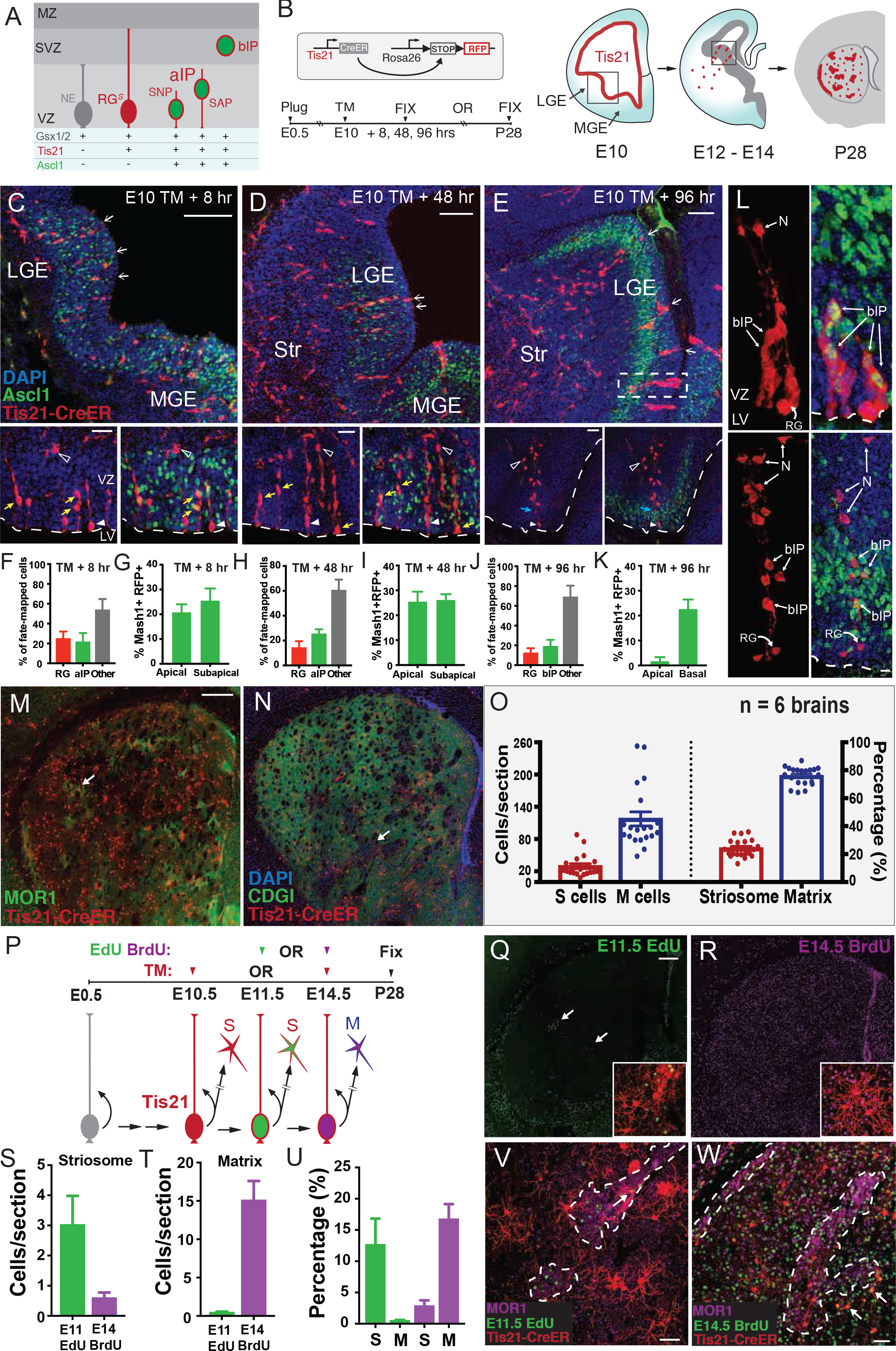
A Radial Glial Cell Lineage in LGE Generates Both Striosomal and Matrix SPNs. (A) Left, schematic representation of LGE progenitor types, including proliferating neuroepithelial cells (NE), self-renewing and neurogenic radial glial cells (RG^*S*^), apical intermediate progenitors (aIPs), and basal intermediate progenitors (bIPs). aIPs further include short neural precursors (SNP) and sub-apical progenitors (SAP). These progenitor types are marked by Gsx1/2, Tis21, Ascl1 expression as indicated. Tis21 marks neurogenic RGs, aIPs and bIPs. IPs co-express Ascl1. VZ: ventricular zone. SVZ: subventricular zone. MZ: mantle zone. (B) Scheme of genetic fate mapping using *Tis21-CreER;Ai14* mice (box). Upon TM induction, Tis21^+^ progenitors are marked throughout the telencephalic VZ as they undergo neurogenesis (E10, schematic). Newborn neurons subsequently migrate to the striatum and self-renewing RGs remain in the VZ (E12-E14, schematic). Fate-mapped SPNs populate both the striosome (red patches) and matrix (surrounding red dots) compartments of striatum (P28, schematic). (C) Eight-hour pulse-chase analysis of Tis21^+^ progenitors (RFP, arrows in upper panel) in LGE (boxed region in B) by TM induction at E10.5 with co-immunostaining for Ascl1 (green) and DAPI counterstain (blue). Bottom panels depict RGs (Ascl1^−^, arrowhead) with radial processes, aIPs (putative SNP/SAPs) with soma located away from the lateral ventricle (Ascl1^+^, yellow arrow), and postmitotic neurons in the emerging MZ (open arrowhead). (D) Forty-eight-hour pulse-chase of Tis21^+^ progenitors (arrows) labeled by TM induction at E10.5 with co-immunostaining for Ascl1. Bottom panels show that self-renewing RGs (Ascl1^−^, arrowhead) remain in VZ at E12.5. aIPs (including a putative Ascl1^+^ SNP; yellow arrows), and postmitotic neurons migrating to the MZ (open arrowhead). (E) Ninety-six-hour pulse-chase analysis of Tis21^+^ progenitors (arrows) labeled by TM induction at E10.5, showing isolated columnar clusters of progenitors and newborn neurons in the LGE proliferative zones at E14. Bottom panels show a RG at the ventricle surface with a long radial fiber and associated Ascl1^+^ bIP in the SVZ (blue arrows). Postmitotic neurons reaching the developing striatum are marked with an open arrowhead. (F, H and J) Percentage of fate-mapped RGs (Ascl1^−^) versus aIPs (Ascl1^+^) at 8, 48 and 96 hours following E10.5 TM induction. “Other” represents cells that could not be readily discerned as RG or aIP. Note that at 96 hours, Ascl1^+^ cells were exclusively localized to the SVZ and thus were scored as bIPs. (G, I and K) Distribution of Ascl1^+^RFP^+^ progenitors within the LGE germinal zone at 8, 48 and 96 hours following E10.5 TM induction. Apical progenitors (soma at ventricle surface), SAPs (soma away from ventricle surface but having apical end-feet), and basal progenitors were scored. (L) Two examples of E10.5 fate-mapped isolated proliferative clusters shown at E14.5 consisting of a putative RG with a radial process running through intermingled Ascl1^+^ bIPs and migrating postmitotic neurons (N) in the SVZ. Top cluster correlates with the dashed box in (E). (M and N) RFP-labeled SPNs fate-mapped from E10.5 Tis21^+^ RGs distribute broadly to both the striosome (arrow; co-labeling with MOR1 in M) and matrix (co-labeled with CDGI in N) compartments. (O) Number (left Y axis) and percentage (right Y axis) of fate-mapped SPNs localized to striosomes and matrix at P28 following TM induction of *Tis21-CreER;Ai14* mice at E10.5. Dots represent individual 50 µm sections selected at 6 representative rostral caudal levels of 6 brains. Error bars = standard error of measurement (SEM). (P) Scheme of combined fate-mapping and birth dating in *Tis21-CreER;Ai14* mice using sequential nucleotide injections of EdU (E11.5, green), followed by BrdU (E14.5, magenta) after E10.5 TM induction. Brains were harvested at P28 to identify co-labeling of RFP with either EdU or BrdU. (Q) Distribution of EdU^+^ cells in P28 striatum following E11 injection; arrows mark clusters of SPNs in striosomes. Inset shows fate-mapped neurons (RFP^+^) co-labeled by E11 EdU injection (green). (R) Distribution of BrdU^+^ cells in P28 striatum following E14 injection, showing broad distribution of labeled cells throughout striatum. Inset shows fate-mapped SPNs (RFP^+^) co-labeled by E14 BrdU injection (magenta). (S, T) Quantification of the numbers of fate-mapped cells per section that incorporated E11 EdU or E14.5 BrdU within striosomes (S) and matrix (M), respectively. The vast majority of E11 EdU^+^ fate-mapped SPNs adopt S cell fate, while the vast majority of E14.5 BrdU^+^ fate-mapped SPNs adopt M cell fate (n = 3 brains from 2 litters). Scale bar: 50 µm (U) Percentage of fate-mapped S versus M cells that co-incorporated E11.5 EdU and E14.5 BrdU, respectively. (V) E10.5 fate-mapped (RFP +) and E11 born (EdU-labeled, green) SPNs distribute to striosomes marked by MOR1 (magenta, dotted line boundary). (W) E10.5 fate-mapped (RFP^+^) and E14 born SPNs (BrdU^+^ pseudocolored green) distribute to matrix (arrows) outside of striosomes marked by MOR1 (magenta, dotted line boundary). Scale bars: 100 µm (top), 20 µm (bottom) in C, D, E; 300 µm in M, Q; 50 µm in V.

To determine the identity of SPNs derived from early Tis21^+^ RGs, we assayed the striatum on postnatal day (P) 28 (Figures 1M–1O). Both striosomal (S) and matrix (M) SPNs were generated, roughly in a 1:4 ratio (Figure 1O). To further probe the lineage relationship between S and M cells, we combined genetic fate mapping of RGs at E10 and consecutive birth dating by EdU at E11 and BrdU at E14. We found that RFP^+^ S cells were predominantly double-labeled by EdU at E11, whereas RFP^+^ M cells were predominantly double-labeled by BrdU at E14 (Figures 1P–1W). Together, these results suggest that along the lineage progression, the same early Tis21^+^ RG population first generates S cells and then M cells, likely via producing several types of IPs.

### A Set of Early Apical Intermediate Progenitors Are Fate-Restricted to Generate Striosomal SPNs

The proneural protein ASCL1 is restricted to more differentiated neurogenic progenitors rather than self-renewing RGs and is important in coordinating the balance between neural progenitor proliferation and cell cycle exit toward neurogenesis (Casarosa et al., 1999; Guillemot and Hassan, 2017). ASCL1 expression exhibits two distinct dynamic patterns: an oscillatory pattern of relatively low ASCL1 levels correlating with cell-cycle progression and proliferation, versus a sustained expression at higher levels driving neurogenic cell division and neuronal differentiation (Imayoshi et al., 2013; Imayoshi and Kageyama, 2014). Thus the *Ascl1-CreER* driver is an effective tool for selectively labeling progenitors biased towards adopting a neurogenic path (Sudarov et al., 2011; Figures 2A–2D). Indeed, throughout E10 LGE, we detected both low and high levels of ASCL1 immunoreactivity in cells within the VZ, suggesting intermingling of cells in various proliferative (oscillatory expression) and neurogenic (sustained expression) states (Figures 2G, 2H, S2A and S2B). We used relatively low-dose TM induction to achieve sparse labeling and to target progenitors with high-level Ascl1 expression, which we verified by co-labeling with anti-ASCL1 antibody (Figure 2H). A brief 8-hour pulse-chase labeling with TM induction in E10.5 *Ascl1-CreER*;*Ai14* embryos reliably labeled aIPs with end-feet at the ventricle surface, with or without basal radial fibers (Figure 2D). Importantly, 18 and 24 hours following induction at E10.5, few or no RFP^+^ progenitors remained in the VZ, in sharp contrast to Tis21^+^ RGs (Figure 1D), indicating that E10 Ascl1^+^ progenitors are aIPs, as opposed to RGs endowed with extensive capacity for self-renewal. Ascl1^+^ aIPs did not express multi-potency markers such as Sox2 (Figure S2C) or the postmitotic neuron marker BIII-tubulin 1(TuJ1; Figures S2C and S2D). It is possible that a lower level of Ascl1 might have been expressed in RGs as well as in aIPs, which may not have been captured by low-dose TM induction. Newly postmitotic RFP^+^ cells expressed typical SPN markers such as CTIP2 as they reached the striatum, but not Nkx2.1, a marker of GABAergic interneurons derived from the medial ganglionic eminence (MGE), suggesting that all fate-mapped cells were SPNs (Figures S2ES2H).

**Figure 2:**
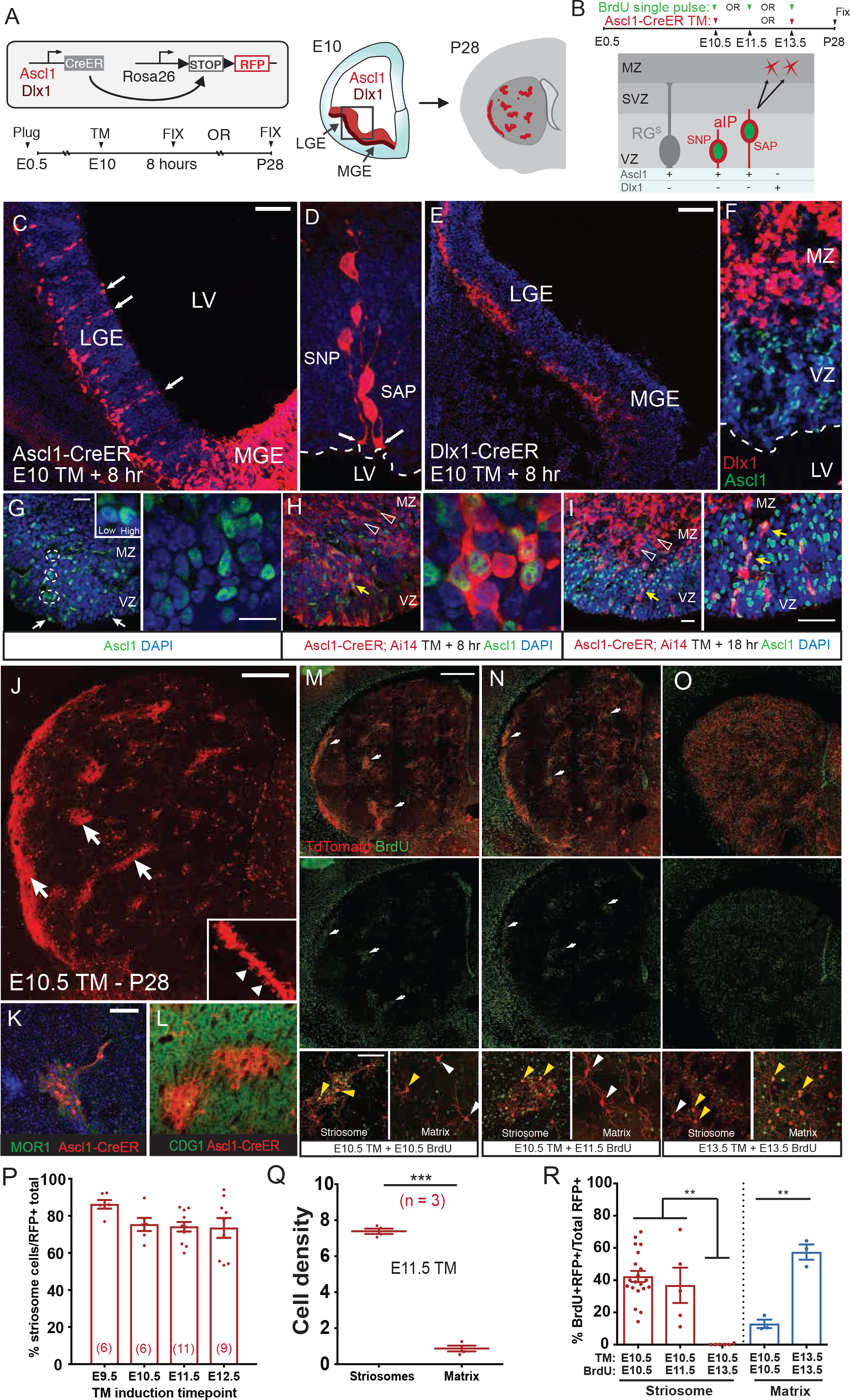
Early Ascl1^+^ aIPs Have Limited Proliferation and Are Fate-Restricted to Produce Striosomal SPNs. (A) Scheme of short- and long-pulse genetic fate mapping by TM induction in *Ascl1-CreER;Ai14* or *Dlx1-CreER;Ai14* mice. Ascl1 (red) and Dlx1 expression (darker red) in LGE are depicted in a schematic E10.5 coronal section. Striosomes in the striatum (red patches) are depicted in a schematic P28 coronal brain section. (B) Scheme of fate mapping aIPs (SNP, SAP) with E10.5 TM induction followed by birth dating with BrdU pulse at E10.5, E11.5, or E13.5 in *Ascl1-CreER;Ai14* mice. The distribution of BrdU^+^ and RFP^+^ cells was assayed in P28 striatum. Differential expression of ASCL1 and DLX1 in RG, aIPs and postmitotic neurons are depicted in the lower panel. (C) Eight-hour pulse-chase of Ascl1^+^ progenitors (arrows) labeled by TM induction at E10.5, arrows indicate aIP with end-feet on the lateral ventricle. Image was taken from the boxed region of the LGE in (A). (D) High magnification view of E10.5 aIPs extending end-feet (arrow) to the ventricle surface (dashed line) with short (putative SAP) or no (putative SNP) vertical radial fibers (arrowheads). (E and F) 8 hour pulse-chase of Dlx1^+^ cells (arrows) labeled by E10.5 TM induction in *Dlx1-CreER;Ai14* embryo. No progenitors in VZ were labeled; instead, postmitotic neurons in MZ were prominently labeled. (G) Ascl1^+^ cells are distributed in a salt and pepper pattern throughout the E10 LGE VZ, with pairs of neighboring progenitors exhibiting high and low levels suggestive of oscillatory expression (inset, white circles), including pairs bordering the lateral ventricle (arrows) and in the VZ but not in the MZ. Scale bars: 20 µm (left), 10 µm (right). See also Figure S2. (H) Eight-hour pulse-chase in *Ascl1-CreER;Ai14* embryo at E10.5 selectively captures cells with high-level ASCL1 in LGE. Many RFP^+^ progenitors in the VZ are actively neurogenic (yellow arrows), while postmitotic neurons (RFP^+^, arrowheads) cease Ascl1 expression and migrate to the MZ. (I) Eighteen-hour pulse-chase shows that most Ascl1^+^ aIPs completed their neurogenic cycles in less than 18 hours, with their postmitotic progenies moving out of the VZ. Only a very small number of Ascl1^+^ RFP^+^ aIPs remain in VZ (arrows). (J and K) SPNs fate-mapped from E10.5 Ascl1^+^ aIPs are destined for striosomes (arrow) in the P28 striatum (dendritic spines in inset). These SPNs are co-labeled by the striosome marker MOR1 (green in J) but not the matrix marker CDGI (green in K). 30 (M and N) SPNs fate-mapped from E10.5 Ascl1^+^ aIPs birth-dated by BrdU injection (green) on E10.5 (M) or E11.5 (N). In each panel, images show BrdU/RFP co-labeling of striosome SPNs at low magnification (arrows, top), BrdU only (middle), and high magnification of co-labeled S cells (bottom left) versus RFP^+^/BrdU^−^ M cells (bottom right). (O) SPNs fate-mapped and birth dated from E13.5 Ascl1^+^ progenitors. Images show BrdU/RFP co-labeling in striatum (top), BrdU only (middle), and high magnification of co-labeled RFP^+^/BrdU^+^ M cells and S cells. (P) Percentage of fate-mapped SPNs localized in striosomes following TM induction at indicated embryonic days. Dots represent individual mice, and their numbers are in parentheses. Error bars, SEM. (Q) Average cell density within the striosome versus the matrix compartment (# of cells/percent compartment area) following E11.5 TM induction. ***p < 0.0001. (R) Quantification of BrdU birth dating in fate-mapped striosome and matrix population, showing the rate of BrdU incorporation at E10.5, E11.5 or E13.5 in S cells (red bars, labeled by E10.5 TM), relative to incorporation of BrdU in M cells at E10.5 or E13.5 (blue bars). Error bars, SEM; **p < 0.001; ***p < 0.0001. Note the absence of S cell production by E13.5. Scale bars: 50 µm in C, E; 20 µm (left), 10 µm (right) in G, I; 300 µm in J, top of M; 100 µm in bottom of M, K.

We then examined the types of mature SPNs in P28 striatum derived from E10.5-labeled Ascl1^+^ aIPs (Figures 2J–2N). Strikingly, early Ascl1^+^ aIPs gave rise almost exclusively to SPNs located within striosomes. These neurons exhibited spiny dendrites and clustered into relatively discrete, widely distributed zones that were positive for the striosome marker mu opioid receptor 1 (MOR1) and negative for the matrix marker CalDAG-GEFI (CDGI) (Figures 2J–2L; Supplemental Movies 2 and 3) (Kawasaki et al., 1998). Both confocal microscopy (Figures 4A and 4C) and serial two-photon tomography (STP; Figure S7; Supplemental Movies 2-4) of the labeled brains demonstrated their morphology and showed that these early aIP-derived SPNs (herein designated as S cells) projected their axons to the dopamine-containing domain of the substantia nigra pars compacta (SNc), a defining feature of striosomal SPNs (Gerfen, 1984; Jiménez-Castellanos and Graybiel, 1989; Crittenden et al 2016). The small fraction of SPNs located in matrix might correspond to the cells described as “exo-patch SPNs” (Smith et al., 2016), long seen in most strisomal preparations (Graybiel and Hickey, 1982; Newman et al., 2015; Graybiel, 1984), likely cells that have not fully completed migration to striosomes but are S cell in type, as the axons of labeled SPNs formed dense projections to SNc, characteristic of S cells. Indeed, throughout the period from E9.5 to E12.5, these *Ascl1*^+^ aIPs almost exclusively generated S cells and not M cells (Figures 2P, 2Q, 5 and S4B).

To further verify that S cells were indeed born from Ascl1^+^ aIPs within this time-window, we combined TM induction at E10.5 followed with BrdU birth dating of their neuronal progeny at several time points (in different litters), including at 4, 24, and 48 hours after TM induction (Figure 2R). These experiments demonstrated that the peak of S cell production from E10.5 aIPs was between E10.5 and E11.5, followed by a sharp decline before E13.5 (Figure 2R). Together, these findings suggest that the early cohorts of Ascl1^+^ aIPs were fate-restricted to produce striosomal SPNs from E10 to ~E12, and confirmed that early Ascl1^+^ aIPs do not linger in the VZ for >24 hours.

As the homeobox gene *Dlx1/2* is also implicated in LGE neurogenesis and striatal development (Anderson et al., 1997), we performed similar fate mapping using the *Dlx1-CreER* driver (Taniguchi et al., 2011). Surprisingly, we found that during the early phase of LGE neurogenesis (E10.5), Dlx1 was not expressed in progenitors in VZ but almost exclusively confined to postmitotic neurons in the mantle zone (MZ; Figures 2E and 2F). Fate mapping of these early-born (E9.5-E12.5) *Dlx1*^+^ neurons to the mature brain showed that they were almost exclusively S cells, similar to those deriving from Ascl1^+^ aIPs during the same embryonic time (also see Figure 5). This result suggests that Dlx1 likely acts down-stream and/or subsequent to Ascl1 during this early phase, possibly in regulating the migration and maturation of postmitotic SPNs.

### A Set of Basal Intermediate Progenitors Are Fate-Restricted to Generate Matrix SPNs

The nearly exclusive fate restriction of early aIPs for S cell production prompted us to examine later phases of LGE neurogenesis from different types of progenitors. At E14.5, 8 hour pulse-chase in *Tis21-CreER;Ai14* embryos continued to label VZ RGs with apical end-feet (Figures 3A and 3E). In addition, large numbers of Ascl1^+^ bIPs were labeled in the SVZ (Figures 3A and 3F), which were frequently engaged in symmetric cytokinesis (Figure 3G), suggestive of neurogenic cell division. Among the total RFP labeled cells, ~5% were putative RGs (located in VZ with end-feet and expression of the proliferation marker Sox2), ~70% were bIPs (Ascl1^+^ in SVZ with no end-feet); and the remaining ~25% of Ascl1^−^ cells in SVZ likely represented other types of bIPs (e.g., Dlx1^+^) as well as new-born postmitotic neurons (Figures 3H and 3I). Notably, in addition to RGs, 24-hour and 72-hour pulse-chase showed the generation of almost exclusively bIPs and not of aIPs (Figures 3B–3D and S3A). In particular, 72-hour pulse-chase demonstrated that Tis21^+^ RGs continued multiple rounds of self-renewal during the late stage of embryonic development with ongoing generation of Ascl1^+^ progenitors in SVZ (Figures 3C, 3D and S3B). These results suggest that during the late phase of LGE neurogenesis (from E14.5 on), Tis21^+^ RGs persist to the end of neurogenesis (Figures S3E and S3F) and begin to predominantly generate a new population of bIPs, possibly in addition to production of a much smaller number of aIPs (see below).

**Figure 3:**
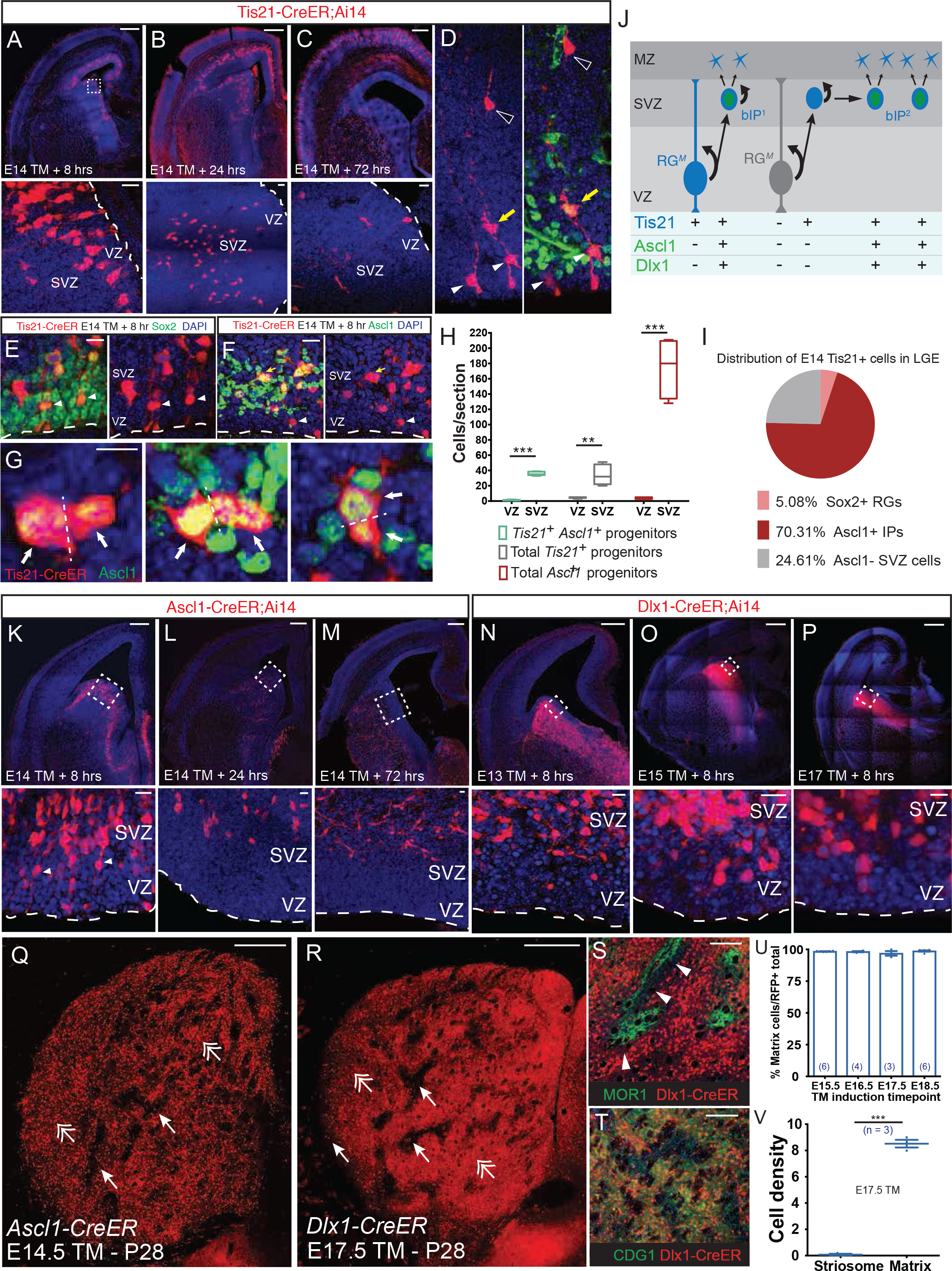
LGE RGs during the Late Phase of Lineage Progression Give Rise to Ascl1^+^/Dlx1^+^ bIPs Committed to the Generation of Matrix SPNs. (A-C) Pulse-chase of E14.5 Tis21^+^ LGE RGs and IPs at 8 (A), 24 (B), and 72 (C) hours in *Tis21-CreER;Ai14* embryos. While post-mitotic SPNs migrate into the striatum, self-renewing RGs (arrowheads) remain in the VZ for at least 72 hours following E14.5 TM induction (bottom panels). (D) Example of a continually self-renewing RG in the VZ (arrowhead) 72 hours following fate-mapping from E14.5 Tis21^+^ RGs. Note the Ascl1^+^ bIP likely derived from the RG (yellow arrow) and the postmitotic progeny neuron migrating to striatum (open arrowhead). (E) Eight-hour TM pulse-chase of Tis21^+^ RGs at E14.5. These RGs were Sox2^+^, Ascl1^−^, and extended end-feet to ventricle surface and radial processes (arrowheads). (F and G) In the same sample as in (E), a large number of bIPs in the SVZ are Ascl1^+^ (F; yellow arrow), many of them in active phases of mitosis (G; dashed lines show plane of cytokinesis), generating putative post-mitotic progenies (arrows). (H) Distribution of Ascl1^+^ IPs in VZ and SVZ following 8-hour pulse-chase in *Tis21-CreER;Ai14* embryo, showing that the vast majority of Tis21^+^, Ascl1^+^ Tis21^+^ and Ascl1^+^ progenitors reside in SVZ at E14, in contrast to the distribution pattern of the same pulse-chase at E10.5 (compare with Figure 1C). (I) Pie chart depicting the percentage of Sox2^+^ RGs, Ascl1^+^ IPs, and Ascl1^−^ SVZ cells among the total population of fate-mapped *Tis21^+^* progenitors at E14. (J) A summary schematic showing that during the late phase of LGE neurogenesis RGs mostly generate bIPs, which may proliferate further and amplify before completing the neurogenic divisions that generate SPNs. Tis21 expression is generally activated in progenitors entering such a neurogenic phase, whether in RG or bIP. (K-M) Pulse-chase of E14.5 Ascl1^+^ IPs at 8 (K), 24 (L), and 72 (M) hours in *Ascl1-CreER;Ai14* embryos. Fate-mapped IPs were completely depleted from VZ by 24 hours after TM induction, and have differentiated into young neurons by 72 hours. Lower panels show magnified view of the boxed areas with VZ and SVZ labeled. (N-P) Eight-hour pulse-chase of Dlx1^+^ IPs at E13 (N), E15 (O) and E17 (P). Lower panels show that Dlx1^+^ IPs did not extend apical or basal processes at any of these stages and thus were exclusively bIPs. Dlx1 is also expressed in postmitotic neurons during this period. (Q) SPNs fate-mapped from E14.5 Ascl1^+^ IPs localized to matrix (double arrows), but not striosomes (arrows) in P28 striatum. (R-T) SPNs fate-mapped from E17.5 Dlx1^+^ IPs localized to matrix (double arrows) but not striosomes (arrows) in P28 striatum (R). These SPNs surround MOR1^+^ striosomes (S) and were co-labeled by the matrix marker CDGI (T), with arrowheads indicating a void in the immediate region surrounding striosomes. (U) Percentage of fate-mapped SPNs that localize to matrix following TM induction to map the output from *Dlx1^+^* IPs at indicated embryonic times. (V) Average cell density within the striosome versus the matrix compartment (# of cells/percent compartment area) following E17.5 TM induction in *Dlx1^+^* IPs. ***p< 0.0001. Scale bars: 300 µm in top of A, B, C, K, L, M, N, O, P; 20 µm in bottom of A, B, C, E, F K, L, M, N, O, P; 5 µm in G; 300 µm in Q, R; 100 µm in S, T.

For a more detailed analysis of the neuronal output of Ascl1^+^ bIPs, we further carried out analogous embryonic fate mapping in E14.5 *Ascl1-CreER;Ai14* mice. Eight-hour pulse-chase of Ascl1^+^ IPs labeled few aIPs and many bIPs, which were identified as progenitors located in the SVZ without apical end-feet (Figure 3K). Importantly, some E14.5-labeled bIPs remained in the SVZ at 24 hours post-induction (Figures 3L S3C and S3D). As the cell-cycle length of LGE IPs at this time is only ~12 hours, this finding corroborated results from live imaging experiments (Pilz et al., 2013) and suggested that some bIPs may serve as transit-amplifying progenitors that proliferate in the SVZ before neurogenesis (Figure 3J). To identify the SPN types derived from late-phase Ascl1^+^ bIPs, we analyzed P28 striatum following E14.5 induction of *Ascl1-CreER;Ai14* mice and detected only M cells (Figures 3Q, 3U and 3V). These results indicate that late-phase *Ascl1*^+^ bIPs are fate-restricted to generate M cells.

As Dlx1 may act downstream and/or in parallel to Ascl1 in the LGE (Long et al., 2009), we also performed systematic fate mapping using the *Dlx1-CreER;Ai14* mice throughout LGE neurogenesis (Figures 3N–3P and 5). Similar to the early-phase *Dlx1^+^* cells’ positions, late phase (E13.5 to 17.5), pulse-chase of Dlx1^+^ cells labeled post-mitotic neurons in the MZ, but excluded RGs and aIPs in the VZ (Figures 3N–3P and S3G-S3J). In striking contrast to the early-phase pulse-chase results, however, late-phase pulse-chase did prominently label bIPs with no apical end-feet throughout the SVZ (Figures 3N–3P). Thus, the recruitment of Dlx1 transcription factor distinguished putative bIPs from aIPs. Furthermore, when assaying fate-mapped SPNs in the mature striatum, we found that late-phase Dlx1^+^ bIPs also specifically produced SPNs of the matrix compartment (Figures 3R–3V). These M cells formed an expansive and dense field of SPNs within which unlabeled striosomes were embedded (Figures 3R–3T). Over 98% of these IP progeny were co-labeled with the matrix marker CDGI, but not with the striosome marker MOR1. These SPNs projected primarily to the non-dopamine-containing SNr, largely sparing the dopaminergic SNc (Figures 4B, 4D and S4C; Supplemental Movie 5). Together with evidence from analysis of Ascl1 and Dlx1 mutant mice (Yun et al., 2002), these results suggest that during the late phase, Dlxl likely acts down-stream and/or in coordination with Ascl1 in bIPs to promote M cell production, in addition to its role in postmitotic SPNs.

**Figure 4:**
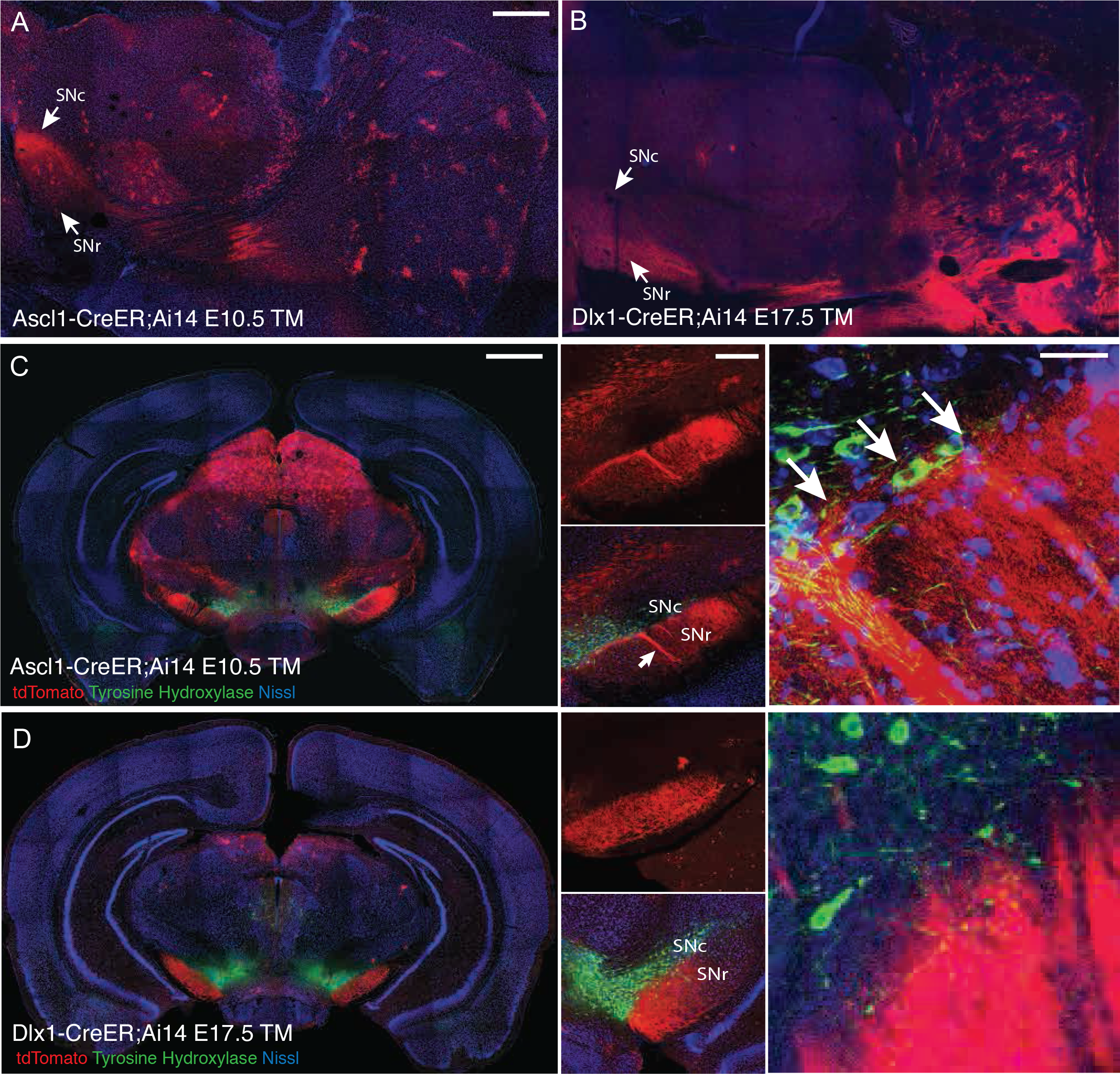
Confirmation of Early Ascl1 Fate-Mapped Striosomal SPNs and Late Dlx1 Fate-Mapped Matrix SPNs by Projection Pattern. (A) Sagittal section showing axon projections from fate-mapped striosomes to substantia nigra following E10.5 TM induction in *Ascl1-CreER;Ai14* mice, terminating densely in the dopaminergic SNc. (B) Sagittal section showing axon projections from fate-mapped matrisomes to substantia nigra following E17.5 TM induction in *Dlx1-CreER;Ai14* mice, primarily terminating in SNr. (C) Coronal section showing axon terminations in substantia nigra from fate-mapped striosomes (E10.5 TM induction in *Ascl1-CreER;Ai14* mice), co-labeled with anti-tyrosine hydroxylase (TH, green) to label dopamine containing neurons in SNc, Middle panels show distribution of striosomal fibers terminating in SNc and SNr (red: tdTomato, green: anti-TH, blue: Nissl), and right panel shows high magnification micrograph of border between SNc and SNr. (D) Coronal section showing axon terminations in substantia nigra from fate-mapped matrix (E17.5 TM induction in *Dlx1-CreER;Ai14* mice), co-labeled with anti- TH (green) to label dopaminergic neurons in the SNc. Middle panels show projections terminate broadly throughout SNr and avoid SNc (red: tdTomato, green: anti-TH, blue: Nissl), and right panel shows high magnification micrograph of border between SNc and SNr. Scale bars: 500 µm in A, C (left); 100 µm C (middle), 10 µm C (right).

### The Sequential Production of Striosomal and Matrix SPNs is Punctuated by a Sharp Transition Period

As Ascl1 and Dlx1 are expressed in the LGE throughout embryonic neurogenesis and are thought to act sequentially as well as in parallel in regulating striatal patterning (Long et al., 2009; Wang et al., 2013), we extended our analyses by performing an extensive set of fate-mapping experiments with Ascl1- and Dlx1*-*driver mice in time-steps of 0.5-1.0 day spanning the entire embryonic neurogenic period from E9.5 to E18.5 (Figures 5 and S4). The results (Figure 5) indicated a surprisingly sharp shift in neurogenic programs for the generation of striosomal and matrix SPNs, likely driven by largely distinct S cell-producing aIPs (aIP^*S*^) and M cell-producing bIPs (bIP^*M*^) pools.

**Figure 5.**
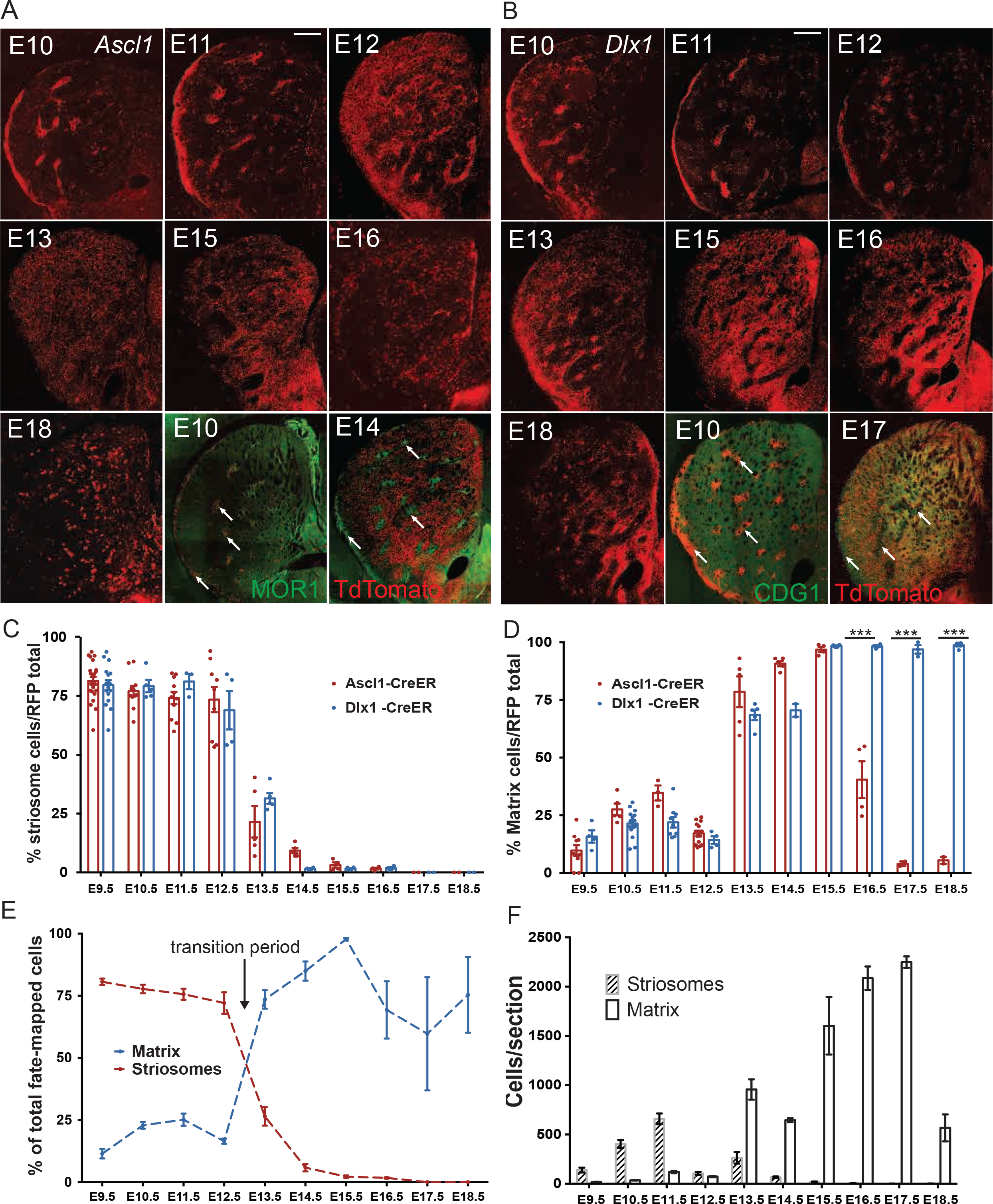
Comprehensive Time-Course of IP-Mediated Neurogenesis Reveals a Rapid Transition from Striosome to Matrix SPN Production. (A) Representative striatum panels of TM induction time points in *Ascl1-CreER;Ai14* mice, including examples of anti-MOR1 co-labeling at peak striosome (E10.5) versus peak matrix (E14.5) cell birth times from *Ascl1^+^* progenitors. (B) Representative striatum panels of TM induction time points in *Dlx1-CreER;Ai14* mice, including examples of anti-CDGI co-labeling at peak striosome (E10.5) versus matrix (E17.5) cell birth times from *Dlx1^+^* precursors. (C) Normalized histogram of striosomal SPN production from *Ascl1*^+^ (red) and *Dlx1*^+^ (blue) progenitors from E9.5 to E18.5. (D) Normalized histogram of matrix SPN production from *Ascl1*^+^ (red) and *Dlx1*^+^ (blue) progenitors from E9.5 to E18.5. (E) Total relative production of striosomes and matrix from *Dlx1*^+^ and *Ascl1*^+^ progenitors pooled from E9.5 to E18.5. (F) Absolute numbers (cells/section) of striosomal or matrix SPNs produced from *Dlx1* and *Ascl1* progenitors (pooled) from E9.5 to E18.5. (n=6-15 sections in 3 mice from 2 litters). Scale bars: 300 µm.

In summary, during an *early phase* (E9.5-E12.5), Ascl1 is expressed in aIPs and Dlx1 in postmitotic neurons; TM induction of both drivers predominantly labeled S cells (Figures 5A and 5C). This early period was followed by a *transition phase* in which there was a rapid decline of S cell production starting around E12.5. From E13.5 on, S cell production became negligible, whereas the labeling of M cells from both drivers was strongly initiated at ~E13.5 and rose sharply within ~24 hours to peak levels. In the subsequent *late phase*, Ascl1 expression shifted from aIPs in the VZ to bIPs in the SVZ, whereas *Dlx1* expression emerged specifically in SVZ bIPs in addition to postmitotic SPNs in the MZ. Between E13.5 and E18.5, the Ascl1 driver initially labeled progenitors producing M cells until ~E16.5, before a sharp decline that was correlated with a switch to the generation of glial cells from the LGE (Figures 5A and 5D). By contrast, the Dlx1 driver persisted in marking progenitors in an active phase of M cell production until the end of neurogenesis at ~E18.5. During both the early and transition phases, the onset and progression of S cell production and the switch from S to M cell production occurred earlier in the Ascl1 driver than in the Dlx1 driver, suggesting that Dlx1 might act downstream of Ascl1 in a transcription cascade that regulates neurogenesis and SPN specification. The MZ expression of Dlx1 in the early and late phase neurogenesis is consistent with this notion (Figures 2E and S3G). The divergence of Ascl1 and Dlx1 cell progeny in the late phase, with Ascl1 driving the production of glia and Dlx1 driving production of M cells, suggests that Ascl1 and Dlx1 probably also act in parallel neurogenetic programs during cell type specification.

Collectively, these results support the existence of two distinct and sequential types of LGE IPs that generate the SPNs of the two major neurochemical compartments of the striatum. An early set of aIP^*S*^ in the VZ generate striosomes with limited neurogenic capacity, whereas a late-activated set of bIP^*M*^ generate matrix SPNs with expanded capacity. Quantification of the total number of fate-mapped S and M cells across the embryonic neurogenic period indicated that the proportion of S and M cells produced was, respectively, approximately 20% and 80% (Figure 5F), matching the approximate ratio of total S and M cells estimated for the mature striatum (Johnston et al., 1990; Mikula et al., 2009).

Given that both Ascl1 and Dlx1 are expressed not only in the LGE but also in the MGE, which generates both cortical and striatal interneurons (Marin et al., 2000; Zhao et al., 2003), it was possible that striatal interneurons were also labeled by TM induction in these driver lines, and that these contributed to, or obscured, the striosome and matrix patterning that we observed. To address the degree to which MGE-derived GABAergic interneurons were intermixed with fate-mapped SPNs derived from the LGE, we devised an intersectional strategy specifically to fate map Ascl1^+^ or Dlx1^+^ progenitors within the MGE (Figure S5). We combined *Ascl1-CreER* or *Dlx1-CreER* drivers with a constitutively active *Nkx2.1-Flp* driver (which targets all *Nkx2*.1 positive MGE progenitors; He et al., 2016) and the intersectional *Ai65* reporter (Madisen et al., 2015) by crossing the corresponding mouse lines. This approach allowed us to restrict fate mapping to *Ascl1^+^* and *Dlx1^+^* progenitors in the MGE and preoptic area, yielding only GABAergic interneurons following TM induction at any given time point (Figure S5). Consistent with previous reports, the overwhelming majority of MGE-derived GABAergic interneurons were born during late embryonic days (~90% from E16.5-E18.5), with very few originating during the period of striosome neurogenesis (Figures S5B-S5D). Similarly, we found parvalbumin-positive and somatostatin-positive interneurons to be scarce within fate-mapped striosomes, but to be located in the matrix compartment when labeled (Figure S6). Cholinergic interneurons were also not observed to be part of Ascl1 and Dlx1 fate-mapped cohorts regardless of induction time, whether from MGE only (intersection) or MGE/LGE (Ai14) (Figure S6). This result suggests that striatal cholinergic interneurons have a distinct progenitor origin from SPNs in the LGE, or that they are generated at a time-point or location not covered by our analysis.

### The Temporal and Spatial Sequence of SPN Settlement in the Striatum Include a Final Cohort That Constitutes Peri-Striosomal Rings

Consistent with previous studies (Krushel et al., 1993; van der Kooy and Fishell, 1987; Newman et al., 2015) Hagimoto et al., 2017; we observed strong spatial gradients in the settling of SPNs along both the lateral-medial and caudal-rostral axes according to the time of Ascl1 and Dlx1expression (Figures 5 and S7; Supplemental Movie 6). More specifically, the SPNs of the matrix compartment not only follow such developmental time-dependent gradient distributions, but are further divided into extended mosaics of cell-clusters (“matrisomes”), which become obvious in tracing experiments demonstrating the input-output organization of the matrix (Flaherty and Graybiel, 1993, 1994; Giménez-Amaya and Graybiel, 1991). Although tract-tracing (Alloway et al., 1999) and 2-deoxy-glucose (Brown et al., 2002) analyses suggest that such patchiness occurs in the striatal matrix in rodents, matrisomes are not visible in conventional histological preparations. Nevertheless, we found a striking non-uniformity in the matrix in our fate-maps: a gap around striosomes with SPNs labeled from E15-E17 cohorts (Figure 3S) that were filled in by a peri-striosomal distribution of the last-born E18 cohort of SPNs (Figure 6). These E18-derived SPNs adhered tightly to the borders striosomes (Figures 6A–6E), and were reminiscent of the peri-striosomal rings demarcated at maturity by immunochemical markers and interneuron distributions observed in primates and humans (Faull et al., 1989; Holt et al., 1997; Graybiel and Ragsdale, 1980). It is thought that these peri-striosomal regions may functionally link striosome and matrix compartments (Banghart et al., 2015; Brimblecombe and Cragg, 2015; Miura et al., 2007). We found that peri-striosomal cells were born almost exclusively during the terminal rounds of neurogenesis at E18.5 (Figure 6). These cells accounted for ~65% of total E18.5 fate-mapped SPNs in whole striatum at P28. Regionally they were more enriched in rostral striatum (~80% of fate-mapped SPNs), reflecting the fact that there were also a significant number of caudal matrix SPNs still being generated at this late time, which mainly populated the medial matrisomes bordering the lateral ventricle (Figures 6C–6F). We also noted similar ring-like structures surrounded MOR1-positive patchy zones in the ventral striatum (Figure 6A). Overall, we conclude that M cells generated from E15.5-E18.5 were deployed in a distal to proximal sequence in relation to the striosomes (Figures 6G–6I). As the RFP-labeled axons of the late-born peri-striosomal M cells innervated both pallidal segments and the SNr, the typical pattern of M cell axons, it is highly likely that at least part of the peri-striosomal M cells correspond to the annular compartment as originally designated by Faull et al. (1989).

**Figure 6:**
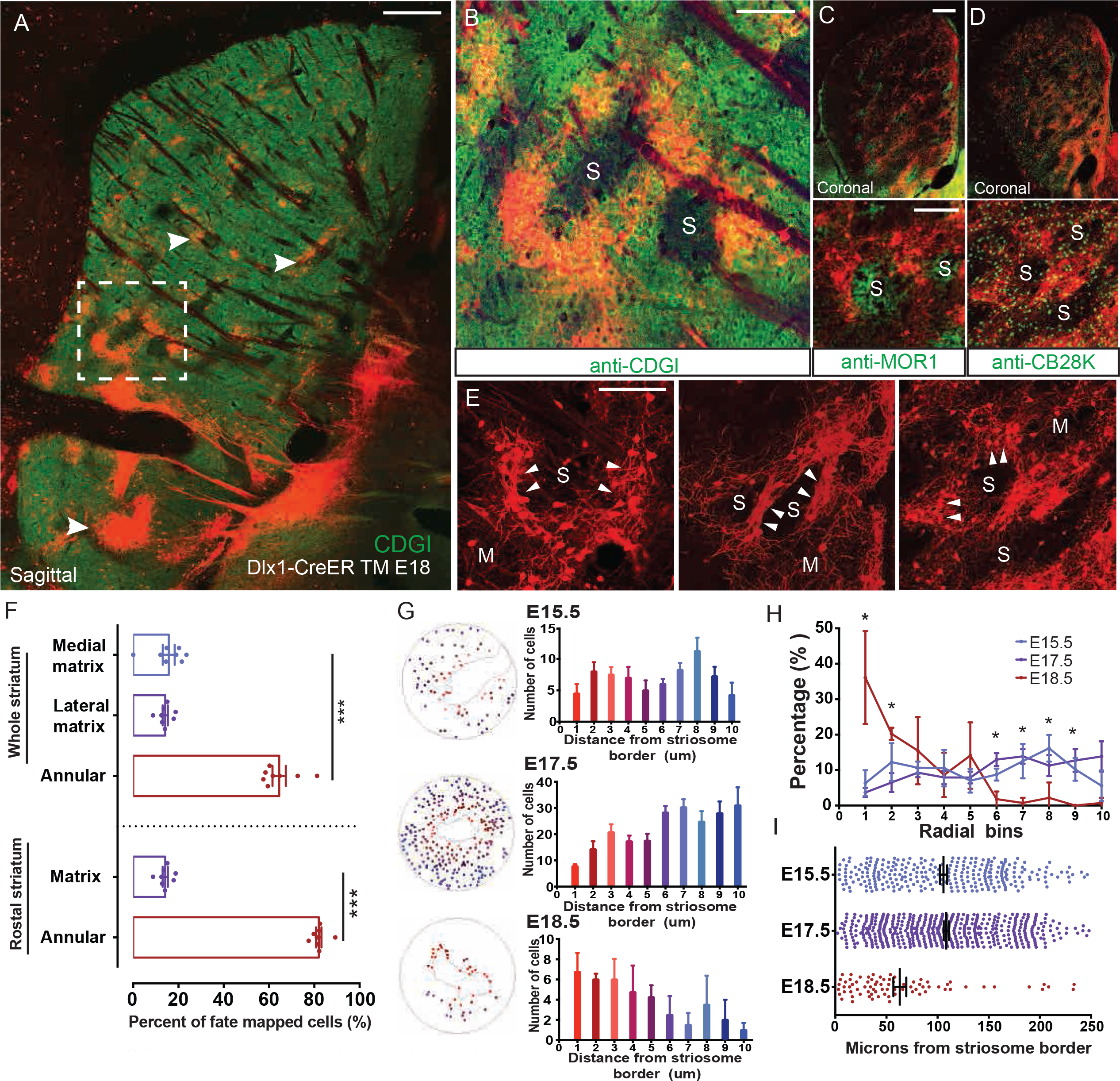
The Annular Sub-Compartment Is Generated through Intermediate Progenitors at the End of the RG^*M*^ Lineage. (A) Sagittal section of P28 *Dlx1-CreER;Ai14* mouse brain following E18.5 TM induction, showing fate-mapped peri-striosomal M cells (arrows) bordering striosomes devoid of CDGI expression (green). (B-D) Rings of peri-striosomal M cells express the matrix markers CDGI (B, rectangle in A) and calbindin D28K (CB28K, D), and adhere tightly to the borders of MOR1^+^ striosomes (C). S: striosomes. (E) Both the neurites and somata of fate-mapped peri-striosomal M cells form dense lattice-like structures (arrowheads) at the interface between striosomes (S) and matrix (M). (F) Fate-mapped SPNs throughout striatum following E18.5 TM induction in *Dlx1-CreER;Ai14* mice. peri-striosomal M cells represent 60-80% of labeled neurons, especially enriched in rostral areas, in addition to a dense band of matrix bordering the lateral ventricle in the medial striatum and sparsely distributed matrix cells throughout the lateral striatum. Error bars, SEM; ***p < 0.0001. (G) Distribution of fate-mapped matrix neurons following E15.5, E16.5 and E18.5 TM induction in *Dlx1-CreER;Ai14* mice, during the onset, peak or closing phase of matrix production, respectively. Cells were grouped into ten 20-µm concentric bins moving radially out from the nearest perpendicular point of a traced MOR1^+^ striosome (red shades closer to and blue shades farther from striosome surface), and the distribution was plotted as a bar graph. n = 4 striosomes from 2 mice per time point. (H) The proportion of total fate-mapped cells in each 20-µm radial bin following E15.5 (purple), E17.5 (blue) or E18.5 (red) TM inductions was plotted for all regions of interest, showing a significant enrichment in cells closer than 40 µm to the striosome border in the E18.5 cohort and a significant enrichment in cells between 120 and 180 µm from striosome borders for both the E15.5 and E17.5 cohorts. n = 4 striosomes from 2 mice per time point. *p < 0.05. (I) Distribution of cells based on distance from striosome borders following E15.5 (purple), E17.5 (blue) or E18.5 (red) TM inductions. Dots represent individual cells, and mean and SEM are indicated in black for each group. Scale bars: 300 µm in A, C (top panel); 100 µm in B, C (bottom panel), and E.

### Direct and Indirect Pathway SPNs in the Striosomes and Matrix Originate Independently within Both aIP^*S*^ and bIP^*M*^

How the neurogenic programs producing striosome-matrix architecture relate to the generation of SPNs of the direct and indirect pathways has long been a major question about basal ganglia organization. Direct and indirect classes of SPNs are distinguished from one another by multiple molecular markers, including their differential expression of D1 or D2 variants of the dopamine receptor (Gerfen et al., 1990; Harrison et al., 1990; Richfield et al., 1989). These markers occur in both striosomes and matrix, but in sharp contrast to the ~1:5 ratio of spatial compartmentalization of striosomal and matrix SPNs, dSPNs and iSPNs are intermixed in both compartments (Fujiyama et al., 2011; Gerfen and Scott Young, 1988; Flaherty and Graybiel, 1994; Banghart et al., 2015; Brimblecombe and Cragg, 2017; Salinas et al., 2016).

To determine whether S- and M-cell fates relate to the acquisition of direct and indirect pathway fates, we distinguished the dSPNs and iSPNs in our genetic fate mapping of the striosome and matrix compartments by relying on the tight correlation between dSPN with D1 receptor expression and iSPN with D2 receptor expression (Gerfen et al., 1990; Harrison et al., 1990; Richfield et al., 1989). We identified these SPNs by fluorescent mRNA double *in situ* hybridization with D1 and tdTomato probes or with D2 and tdTomato probes performed on brain sections in *Ascl1-CreER;Ai14* and *Dlx1-CreER;Ai14* mice induced at specific embryonic times. Thus we simultaneously identified neurons of the striosome and matrix compartments in terms of their D1 and D2 expression levels (Figure 7).

**Figure 7:**
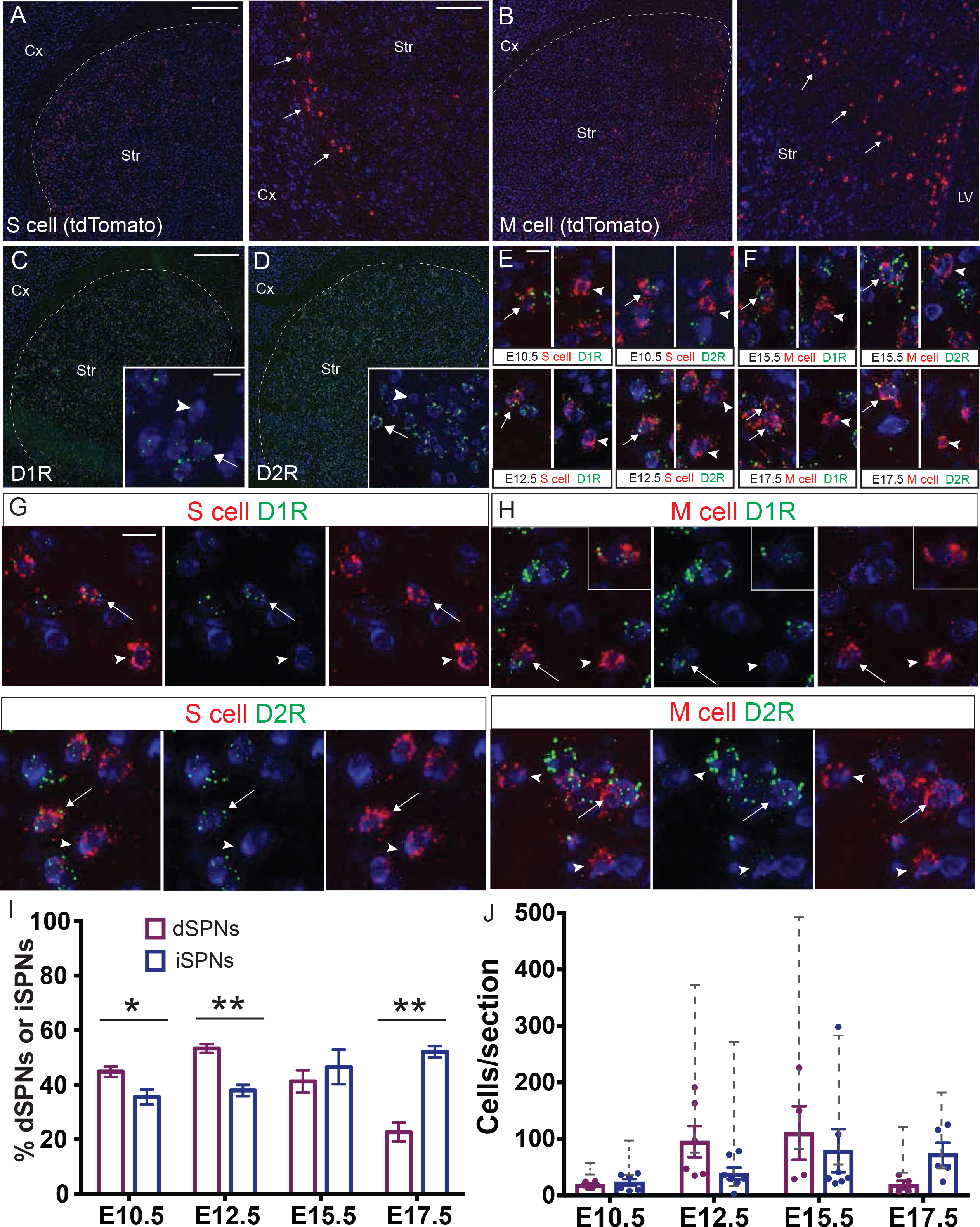
Production and Integration of Direct and Indirect Pathway SPNs within the Striosome-Matrix System. (A) Fluorescent *in situ* hybridization (FISH) signal from a branched-chain DNA (bDNA) probe targeting the mRNA of tdTomato expressed in P28 SPNs following E10.5 TM in *Ascl1-CreER;Ai14* mice matches the signature expression pattern (arrows) of striosome cells along the lateral border of striatum Red: tdTomato mRNA, Blue: Nissl. Scale bars: 300 µm in left panel, 100 µm in right panel. (B) FISH signal for a bDNA probe targeting the mRNA of tdTomato expressed in P28 SPNs following E17.5 TM in *Dlx1-CreER;Ai14* mice matches the signature matrix pattern (arrows) in medial striatum. (C) FISH signal for a bDNA probe targeting the mRNA of the dopamine D1 receptor (D1R) is distributed throughout about 50% of Nissl positive cells in striatum (Green: D1R mRNA, Blue: Nissl, inset: Arrows: positive cell, arrowhead: negative cell). Scale bars: 300 µm,10 µm in inset. (D) FISH signal for a bDNA probe targeting the mRNA of the dopamine D2 receptor (D2R) is distributed throughout about 50% of Nissl positive cells in striatum (Green: D2R mRNA, Blue: Nissl, inset: Arrows: positive cell, arrowhead: negative cell) (E) Representative panels of double *in situ* hybridization marking either D1 or D2 expressing cells in striosomes born at E10.5 (top group) or E12.5 (bottom group). Arrows: co-labeled cells, arrowheads: tdTomato-only cells. Scale bar: 10 µm. (F) Representative panels of double *in situ* hybridization marking either D1 or D2 expressing cells in matrix born at E15.5 (top group) or E17.5 (bottom group). Arrows: co-labeled cells, arrowheads: tdTomato-only cells. (G and H) Differential expression of the D1R or D2R in fate-mapped striosomal (G) and matrix (H) SPNs. Arrows: co-labeled cells, arrowheads: tdTomato-only cells. Scale bar: 10 µm. (I) Proportions of dSPNs and iSPNs within fate-mapped striosome cohorts following E10.5 or E12.5 TM induction and within fate-mapped matrix cohorts following E15.5 or E17.5 TM induction. Error bar: SEM; *p < 0.05, **p & 0.001. (J) Number of dSPNs and iSPNs within fate-mapped striosome cohorts and matrix cohorts (dots represent individual sections, bars project mean values, error bar: SEM). Grey brackets display the range of the number of total RFP^+^ cells per section detected by in situ probe in samples pooled from both high and low TM dose litters.

The spatial pattern of tdTomato mRNA-expressing S cell clusters was readily distinguishable from the M cell distributions. By contrast, D1 and D2 mRNA signals were evenly distributed in apparently equal numbers intermixed throughout the striatum (Figures 7C and 7D). Among S cells labeled by TM induction of *Ascl1-CreER;Ai14* mice at E10.5 and E12.5, approximately equal ratios of D1 and D2 cells were detected, with a slight enrichment of D1 cells (Figure 7I). Similarly, among M cells labeled by TM induction of *Ascl1-CreER;Ai14* mice at E15.5, we found equal numbers of D1 and D2 cells (Figure 7I). Among M cells labeled by TM induction of *Dlx1-CreER;Ai14* mice at E17.5, again both D1 and D2 cells were detected, but with a significant enrichment of D2 cells (Figure 7I). Together, these results indicate that, at least at the population level, the generation of dSPNs vs. iSPNs is not specified by a program analogous to one that generates striosome vs. matrix compartments. The remarkable conclusion suggested by these results is that the specification of a direct-indirect pathway fate appears to be mainly independent of striosome-matrix fate and manifests within both the aIP^*S*^ and bIP^*M*^ progenitor pools, though with a slightly skewed distribution toward a D1-striosome/D2-matrix bias.

## DISCUSSION

Two fundamental categories of vertebrate behaviors that enhance fitness encompass action selection, involving evaluation of multiple options, and action evaluation, based on behavioral outcome. In mammals, action selection and evaluation are crucially regulated by basal ganglia circuitry (Amemori et al., 2011; Graybiel, 2008; Hikosaka et al., 2014), through which the cerebral cortex modulates a wide range of volitional and habitual behaviors. As the recipient of multi-modal information represented in different neocortical areas and thalamic nuclei, the striatum is thought to reconfigure and then to forward the resultant signals to basal ganglia output circuits to initiate and maintain appropriate actions while suppressing others, and to evaluate their outcomes (Mink, 1996; Graybiel and Grafton, 2015). To support such sophisticated operations, the circuit architecture of the striatum is subdivided into striosome and matrix compartments, each further containing direct and indirect pathway subdivisions of SPNs as defined by their selective expression of dopamine receptor variants. Here, we have discovered that the basic plan for assembling the striatal circuit architecture has its root in a developmental program embedded in the LGE RGs and their lineage progression, which generate distinct fate-restricted IPs to specify SPNs into striosome and matrix types, upon which an independent and secondary mechanism divides SPNs into the direct and indirect pathways. This set of findings raises the possibility that the deployment of different progenitor types may reflect an evolutionarily conserved mechanism to assemble striosomal and matrix circuits that mediate distinct behavioral categories involving action selection and evaluation (Grillner and Robertson, 2015; Amemori et al., 2011; Graybiel, 1997; Graybiel and Kimura, 1995; Graybiel, 1997; Houk et al., 1994) while the approximately equal ratio of intermixed direct and indirect pathway SPNs within the two compartments could represent a mechanism for more flexible configuration of functional sub-circuits based on specific outcome evaluation and motor outcome (i.e., experience). This view further raises the intriguing and more general possibility that, as in invertebrates in which cell lineage and developmental genetic program shape circuits and behavior (e.g., Harris et al., 2015), a developmental ground plan in mammalian telencephalic neural progenitors may direct the assembly of certain basic forebrain circuit organization underlying high-order behaviors.

### Lineage Dependent and Independent Production of Different Types of SPNs

The actions of specific transcription factors, such as Gsx1/2, Ascl1, and Dlx1/2, as well as Notch signaling, mediate cell autonomous and non-autonomous regulation of neurogenesis in the LGE and control ordered production of striatal neurons (Mason, 2005; Yun et al., 2002). It has been suggested that the early LGE contains Gsx1/2^+^ neuroepithelial (NE) cells that give rise to multiple progenitor states characterized by Ascl1 and Dlx expression (Martín-Ibáñez et al., 2012; Yun et al., 2002). Ascl1^+^/Dlx1/2^−^ and Ascl1^+^/Dlx1/2^+^ progenitors are inferred to emerge in a sequence (Martín-Ibáñez et al., 2012) and to interact through Notch-mediated lateral inhibition to coordinate proliferation and neurogenesis and to regulate orderly production of early- and late-born SPNs (Mason, 2005). Here, by combining cellular resolution genetic fate mapping and neuronal birth dating, we were able to identify distinct progenitor types, to uncover their lineage progression, and to track their neuronal progenies to SPN types. Based on our findings, we propose a working model of the progenitor and lineage mechanisms underlying the production of SPN types and their influence on striatal circuit architecture (Figure 8).

**Figure 8:**
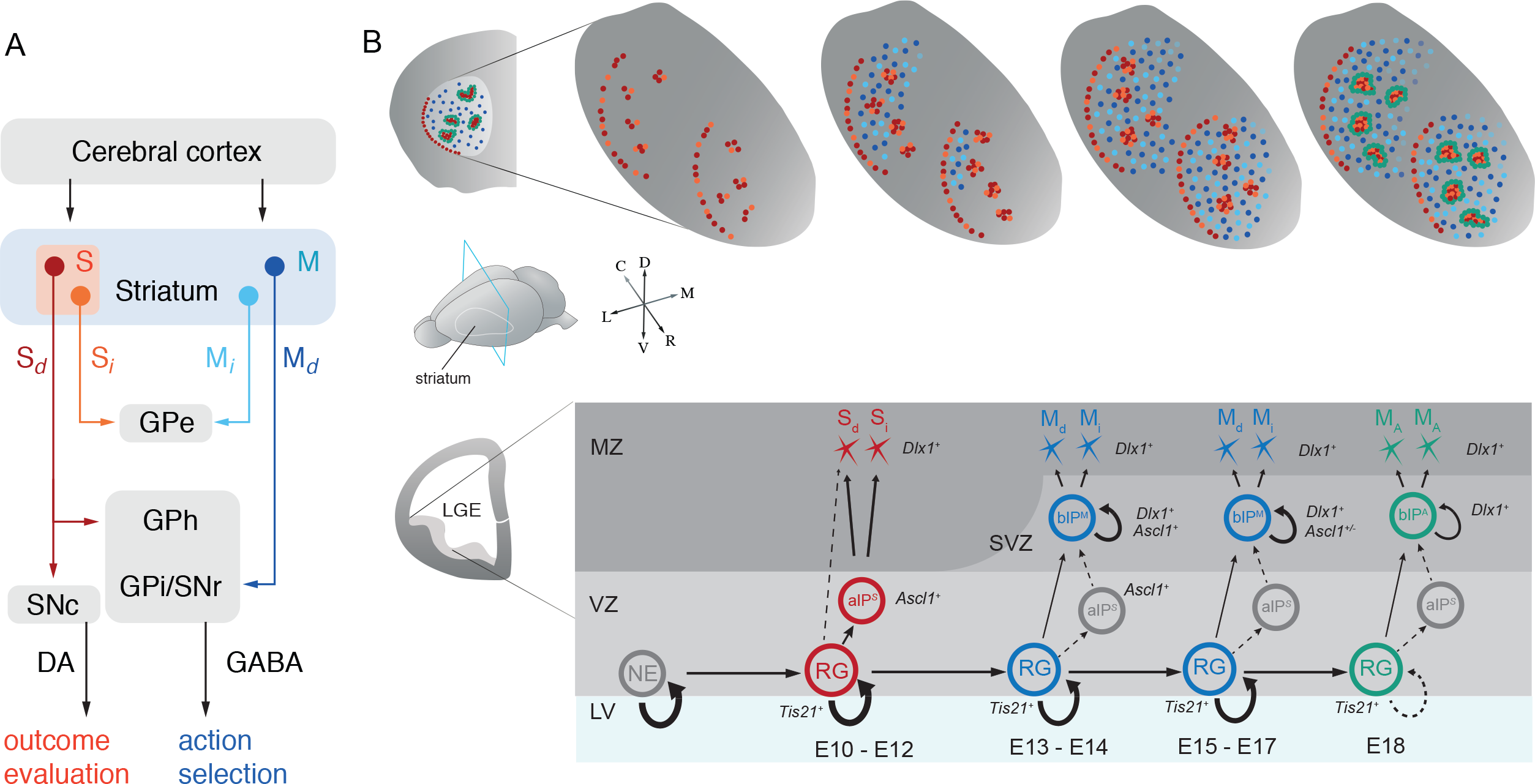
A Model on the Progenitor and Lineage Basis of SPNs that Constitute the Striosome and Matrix Compartments, and Direct and Indirect Pathways. (A) A schematic of the organization of striatal compartments and pathways that mediate different behaviors (GPe: globus pallidus externus, GPi: globus pallidus internus, GPh: globus pallidus habenula, S_d_: striosome direct pathway, S_i_: striosome indirect pathway, M_*d*_: matrix direct pathway, M_*i*_: marix indirect pathway). (B) At the onset of LGE neurogenesis marked by Tis21 expression in radial glial progenitors (RG) ~E10, RGs initiate a lineage program that sequentially generates all striatal projection neurons (SPNs). During an early phase (E10-E12), RGs generate a set of Ascl1^+^ aIP^*S*^s (including SNP and SAPs), which are fate-restricted to generate striosomal SPNs. It is possible that RGs also generate SPNs via direct neurogenesis (dashed line). Both direct (S_d_, light red) and indirect (S_i_, dark red) pathway SPNs are generated from aIP^*S*^s, though we cannot distinguish whether a single aIP^*S*^ generates either or both S_*d*_ and S_*i*_. S cells populate the striatum in a lateral to medial spatial sequence by birth time (top panel). Following a rapid transition period around E13, RGs generate a set of Ascl1^+^ and Dlx1^+^ bIP^*M*^s, which are fate-restricted to generate matrix SPNs. Throughout this late phase (E15-E17), M cells are produced from mostly Dlx1^+^ bIPs, while Ascl1^+^ bIPs taper M cell production and switch to gliogenesis. Again, both direct (M_*d*_, light blue light = D1R^+^) and indirect (M_*i*_, dark blue = D2R^+^) pathway SPNs are generated from bIP^*M*^s; and we cannot distinguish whether a single bIP^*M*^ generates either or both M_*d*_ and M_*i*_. M cells populate the striatum in a caudal to rostral and lateral to medial sequence by birth time (top panel). During the final phase, Dlx1^+^ bIPs produce projection neurons of the peri-striosomal annular compartment (A cells). Thus, M cell generation further manifests a distal to proximal sequence in relation to the striosome. Thus along the RG lineage progression, different progenitor types and cell birth time specify distinct types of striosome and matrix SPNs (S, M and A cells) and shape the spatiotemporal construction of striatal compartments and likely their input-output connectivity.

According to this model, a multi-potent radial glial lineage sequentially allocates distinct intermediate progenitor (IP) types to build striosome-matrix compartmentalization, and the division of direct and indirect pathways occurs within each IP type secondary to the compartmental divide. More specifically, the onset of neurogenesis in the VZ of early LGE (~E10) is marked by Tis21 expression in RGs, which self-renew over multiple rounds to generate several types of IPs and all types of striatal SPNs. In the early phase of this lineage progression, RGs may generate neurons directly and/or through aIPs (including SNPs and SAPs) with dynamic ASCL1 expression patterns. Early aIPs (herein termed aIP^*S*^cells) with sustained high ASCL1 levels are committed to produce striosomal SPNs with no or likely no more than one round of amplification, thus limiting their neurogenic capacity. These aIP^*S*^s produce both striosomal dSPNs and iSPNs in an approximately equal ratio. During a brief transition period starting ~E12.5, RGs undergo a rapid shift from generating aIPs to generating bIPs that are fate-restricted to produce M cells (i.e., bIP^*M*^). Dlx1 is activated downstream and/or in parallel to Ascl1 in bIP^*M*^. A possible rapid depletion of aIP^*S*^s may result in a sharp decline of S cell production, while the rapid rise in the production of bIP^*M*^s promotes M cell genesis. In the next phase (E14.5-E17.5), Ascl1^+^ bIP^*M*^s first promote M cell production and then switch to the generation of glial cells. By contrast, Dlx1^+^ bIP^*M*^s persist in driving highly prolific M cell production throughout the late phase of LGE neurogenesis. Many bIP^*M*^s may undergo one or more rounds of transit-amplifying proliferations before the final neurogenic division, thereby increasing their neurogenic capacity, in addition to the prolific generation of bIPs by RGs through aIPs, and likely proliferation of the RG pool itself. Such mechanisms likely 15 account for the significant numerical expansion of the matrix relative to the striosome compartment. At the final stage (~E18.5), Dlx1^+^ IPs generate SPNs destined to form peri-striosomal ring structures, of which some subset likely contribute to the annular compartment. Throughout bIP^*M*^-mediated neurogenesis, as during the aIP^*S*^-mediated period that precedes it, a similar ratio of matrix dSPNs and iSPNs are generated, with a bias toward iSPNs at the end of bIP^*M*^ neurogenesis.

Our fate-mapping method cannot distinguish direct vs. indirect neurogenesis from Tis21^+^ RGs. Additionally, the mechanism underlying the shift of RG state from aIP*^S^* to bIP^*M*^ production is unknown and might involve RG intrinsic (e.g., tracking cell division number) or extrinsic (e.g., signaling from increasing number of S cells) processes; transcriptome analysis of early and late RGs may uncover the molecular basis.

### The Relation of Striosome-Matrix to Direct-Indirect Pathway SPN Origins

A remarkable finding from our fate-mapping experiments is that dSPNs and iSPNs appear to derive from a different developmental process than that giving rise to the striosome and matrix compartments: both dSPNs and iSPNs derive from both aIP^*S*^ and bIP^*M*^ pools and in approximately equal or biased ratios, depending on the embryonic day, throughout the LGE neurogenic window. How could such a consistent pathway division be set up within the developmental process giving rise to striosomes and matrix? One possibility is that when S or M cells are first specified, they are neutral with respect to the D1-D2 distinction, so that these phenotypes are acquired in response to environmental factors or cell-cell interactions. An alternative possibility is that the D1 and D2 fates emerge together with the fundamental distinction into striosome or matrix identities. Thus, it is formally possible that, within the aIP^*S*^ and bIP^*M*^ pools, subsets of IPs are further fate-restricted to generate direct vs. indirect pathway S cells, and direct vs. indirect pathway M cells. We also note that a stochastic mechanism (Boije et al., 2014) might be well suited to generate the characteristic ratio of the two pathway SPNs, as stochastic decisions can produce a mosaic of fates within a population of cells, and they can be integrated into developmental programs and directed to yield robust and reproducible outcomes (Boije et al., 2014; Johnston and Desplan, 2010). Such a stochastic mechanism for D1 vs. D2 fate could manifest itself either at the step of IP generation from RGs or at the final neurogenic division of individual IPs. Nearly equal or biased ratios of highly intermixed dSPNs and iSPNs generated by stochastic fate choice from IPs could lay out a template in both striosomes and matrix upon which learning and experience can shape behaviorally relevant functional sub-circuits.

### The Significance of Distinct IPs and Their Differential Amplification for the Production of Striosomal and Matrix SPNs

In addition to differences in input-output connectivity and behavioral function, the striosome and matrix compartments differ substantially in size and maintain an approximately 1:4 ratio that is conserved across mammalian species. The mechanisms underlying this size difference between the two compartments is not well understood. Our finding that distinct IP types with different neurogenic capacities are fate restricted for S vs. M cell production provides plausible mechanisms. First, a larger number of bIPs^*M*^ are generated from RGs during a longer late phase and possibly through late-phase aIPs that further amplify bIPs^*M*^ numbers. Second, individual bIPs^*M*^ undergo more rounds of transit amplification compared to early aIPs^*S*^. We suggest that aIP^*S*^-mediated generation of SPNs in the early phase forms an evolutionarily conserved basic template for evaluative circuits. On the other hand, bIP^*M-*^mediated amplification (starting from ~E13) dramatically increases the production of matrix SPNs and could underlie the development of the much larger matrix compartment with potential further diversification such the peri-striosomal rings. The greatly expanded matrix SPN population may provide the basis underlying flexibility for building a large repertoire of basal ganglia circuits contributing to action selection and motor skill learning in parallel with the expansion of the neocortex.

The recognition of the distinction between the developmental plan for striosome-matrix compartmentalization and its relation to the direct-indirect pathway subdivision of SPNs opens the way to a deeper examination of striatal circuitry linking development and behavior. As these two organizational plans are now known to harbor different disease vulnerabilities, for example, of striosomes in mood disorders in Huntington’s disease patients and of D2-bearing striatal neurons in Parkinson’s disease, these findings may have significance for understanding the developmental and circuitry etiology of these clinical disorders (Crittenden and Graybiel, 2016).

## EXPERIMENTAL PROCEDURES

### Mice

Mouse lines are described in the Supplemental Experimental Procedures. All experiments and procedures are performed according to NIH guidelines and approved by the IACUC at Cold Spring Harbor Laboratory and at Massachusetts Institute of Technology.

### Inducible Genetic Fate Mapping

Inducible genetic fate-mapping methods are described in the Supplemental Experimental Procedures.

### Tamoxifen Induction and BrdU Birth Dating

To reduce toxicity to pregnant females and enable sparse cellular labeling, pregnant female mice were submitted to gavage with low to medium dose tamoxifen (dissolved in corn oil by gentle agitation for 24 hours), using an 18-gauge 50 mm gavage tool. Doses were in the range of 0.15 mg/30 g body weight (BW) up to 5 mg/30 g BW (5 mg/Kg BW–165 mg/Kg BW), with most experiments performed using doses below 1.5 mg/30 g BW. For BrdU birth dating, pulses of BrdU were intraperitoneally injected into pregnant mice at a dose of 5 mg/100 g BW, spaced 4 or 24 hours after TM induction on E10.

### Immunohistochemistry and Microscopy

Immunohistochemistry and microscopy are described in the Supplemental Experimental Procedures.

### Double Fluorescent *in situ* Hybridization

Two-probe fluorescent *in situ* hybridization was performed on fresh frozen 12 µm thick cryosections of P28 mouse brains using the QuantiGene ViewRNA ISH assay from Affymetrix. Briefly, tissue was treated with formaldehyde fixation, protease digestion, target and label probe incubation, and buffer washing according to the QuantiGene Affymetrix protocol. Tissue was then co-labeled with fluorescent Nissl and imaged using a DeltaVision epifluorescent microscope (details described in Supplemental Experimental Procedures).

### Imaging and Serial Reconstruction of Whole Brains

Imaging and methods for serial reconstruction of whole brains are described in the Supplemental Experimental Procedures.

### Quantification and Data Analysis

Quantification and data analysis are described in the Supplemental Experimental Procedures.

## AUTHOR CONTRIBUTIONS

Z.J.H and A.M.G. conceived and organized the study. S.M.K. and Z.J.H. designed the experiments. S.M.K., P.W. and M.H. designed and generated Tis21-2A-CreER and Nkx2.1-ires-Flp knock-in mouse lines. S.M.K and R.R. conducted primary genetic fate-mapping experiments and embryonic pulse-chase experiments. J.L., L.G.G. and A.M.G. performed comprehensive tamoxifen inductions and histology for each half-day fate-mapping time point and collected brain samples for *in situ* hybridization experiments. S.M.K. completed fluorescent double *in situ* experiments. S.M.K., Y.K. and P.O. performed serial two-photon imaging experiments. J.L. and K.M. contributed to immunostaining and imaging of striatal interneuron markers in fate-mapped cells. Z.J.H, A.M.G. and S.M.K. wrote the manuscript.

## ACKNOWLEDGMENTS

We are grateful to Sang Yong Kim for help with generation of knock-in mice, to Nour El-Amine for help with imaging of fluorescent double *in situ* samples, and to Gord Fishell, Songhai Shi, Marcus Stepheson-Jones, Bo Li, and Yasuo Kubota for discussions and invaluable comments on early versions of this manuscript. This work was supported in part by a grant from the Simons Foundation to Z.J.H. and A.M.G., by CSHL Robertson Neuroscience Fund to Z.J.H., by a grant from the CHDI Foundation (A-5552) to A.M.G., and by support from the Saks Kavanaugh Foundation to A.M.G. S.M.K. was supported by NRSA F30 Medical Scientist Predoctoral Fellowship 5F30MH102002-03, and L.G.G. was supported in part by a Postdoctoral Fellowship from the William N. & Bernice E. Bumpus Foundation. Work is also Supported also by NIMH grant 1S10OD021759-01

## SUPPLEMENTAL EXPERIMENTAL PROCEDURES

### Mice

All mouse colonies at Cold Spring Harbor Laboratory (CSHL) were maintained in accordance with protocols approved by the IACUC at CSHL. The following mouse strains were used for fate-mapping experiments: Ascl1-CreER (Kim et al., 2011), Dlx1-CreER (Taniguchi et al., 2011), Nkx2.1-ires-FLP-O (He et al., 2016), Ai14 (Madisen et al., 2010), Ai65 (Madisen et al., 2015), and Tis21-2A-CreER (unpublished). A gene targeting vector for Tis21-2A-CreER was generated using a PCR-based cloning approach as described before (Taniguchi et al., 2011) to insert a 2A-CreER construct immediately after the STOP codon of the Tis21 gene. The targeting vector was linearized and transfected into a 129SVj/B6 F1 hybrid ES cell line (V6.5, Open Biosystems). G418-resistant ES clones were first screened by PCR and then confirmed by Southern blotting using probes against the 5’ and 3’ homology arms of the targeted site. The loxP-Neo-loxP cassette in _the_ founder line was removed by breeding with CMV-cre transgenic mice (JAX 006054).

Once generated at CSHL, Ascl1-CreER and Dlx1-CreER mice were sent to the Massachusetts Institute of Technology (MIT), rederived and maintained under a standard light/dark cycle with free access to food and water in MIT housing facilities. All experimental procedures were approved by the Committee on Animal Care at MIT and were carried out in accordance with the U.S. National Research Council Guide for the Care and Use of Laboratory Animals.

### Inducible Genetic Fate Mapping

Genetic fate-mapping experiments were carried out in parallel at CSHL and MIT. Inducible fate-mapping experiments were conducted by crossing either Dlx1-CreER or Ascl1-CreER mice to the Cre-dependent Ai14 reporter. Intersectional inducible fate-mapping experiments were conducted by crossing Nkx2.1-ires-FLP-O;Ai65 double transgenic mice to a CreER driver line (backcrossed 6 generations to Swiss-Webster background). Upon observing a vaginal plug, with the noon of that day counted as E0.5, tamoxifen was administered by gavage to the pregnant female at a desired embryonic time point (Taniguchi et al., 2011). Adult progeny mice with appropriate genotypes were anesthetized with 2.5% avertin before transcardial perfusion with 4% paraformaldehyde (PFA) onP28. Brains were post-fixed in 4% PFA for 12 hours. In 8-hour pulse-chase experiments, E10–E13 embryos were lightly fixed by immersion in 4% PFA starting 8 hours after TM induction and frozen for cryosectioning after overnight cryoprotection in 30% sucrose. Later stage embryos (E14–E18) were perfused with 4% PFA using a 30-gauge needle and syringe before being post-fixing brains for 6 hours and embedding in 1.8% agarose for vibratome sectioning (Taniguchi et al., 2011).

### Tamoxifen Induction and BrdU Birth-Dating

To reduce toxicity to pregnant females and enable sparse cellular labeling, pregnant female mice were gavaged with low to medium dose tamoxifen (dissolved in corn oil by gentle agitation for 24 hours), using an 18-gauge 50-mm gavage tool. Doses were in the range of 0.15 mg/30 g BW up to 5 mg/30 g BW (5 mg/Kg BW–165 mg/Kg BW), with most experiments at CSHL performed using doses below 1.5 mg/30 g BW and all experiments at MIT performed using doses of 3 mg/30 g BW. For BrdU birth-dating, pulses of BrdU were intraperitoneally injected into pregnant mice at a dose of 5 mg/100 g BW, spaced 4 hours or 24 hours after TM induction on E10, similar to previously described (Taniguchi et al., 2011).

### Immunohistochemistry and Microscopy

Fixed adult brains were serially sectioned at 55 µm using a vibratome and collected into well plates. Sections were blocked in 10% normal goat serum and 0.1% triton-X100 in 1XPBS, then incubated in primary antibodies (diluted in blocking solution) at 4°C overnight: anti-RFP (rabbit polyclonal antibody, Rockland Pharmaceuticals), anti-MOR1 (rabbit polyclonal, immunostar), anti-Calbindin 28K (rabbit polyclonal, Millipore), anti-CDG1 (rabbit polyclonal, Crittenden et al., 2009), anti-Tyrosine Hydroxylase (rabbit polyclonal, Millipore), anti-Ascl1/Mash1 (mouse monoclonal, BD biosciences), anti-BrdU (rat monoclonal Accurate Chemical), anti-TuJ1 (mouse monoclonal, Covance), anti-Parvalbumin (mouse monoclonal, Sigma), anti-Somatostatin (rabbit polyclonal, Millipore), anti-ChAT (rabbit polyclonal, Santa Cruz biotechnology). Sections were then rinsed and incubated with appropriate Alexa Fluor dye-conjugated IgG secondary antibodies (1:800 goat anti-rabbit/mouse/rat, Life Technologies) and mounted in Fluoromount-G (SouthernBiotech). Sections were counterstained with NeuroTrace Nissl Stain (Molecular Probes) or DAPI. For BrdU immunostaining, sections were denatured in HCl (2 N) at 37°C for 45 min, then neutralized in sodium borohydrate buffer (0.1 M pH 8.5) twice for 10 min at room temperature before the normal staining protocol.

At MIT, 40-um thick coronal frozen sections through the striatum were cut with a sliding microtome and were stored in a solution of 0.1M NaKPO_4_ and 0.4% Na azide until use. For immunofluorescence immunohistochemistry, sections were rinsed in 3x2 min in 0.01M phosphate buffered saline (PBS) containing 0.2% Triton-X-1000 (Tx) and were blocked in tyramide signal amplification (TSA) blocking 2 reagent (Perkin Elmer) for 20 min. The sections were then incubated with primary antibodies in TSA blocking reagent in PBS-Tx for 24 hours at 4°C with gentle shaking: anti-RFP (rabbit/mouse polyclonal/monoclonal, Rockland Pharmaceuticals), anti-MOR1 (rabbit polyclonal, Immunostar/Abcam), anti-CDG1 (rabbit polyclonal, Graybiel laboratory), anti-parvalbumin (mouse monoclonal, Millipore), anti-somatostatin (rabbit polyclonal, Millpore), and anti-ChAT (rabbit polyclonal, Millipore). After 24-hour incubation, sections were rinsed 3x2 min in PBS-Tx and incubated for 2 hours in secondary antibody solution with the appropriate Alexa Fluor dye-conjugated IgG secondary antibodies (1:300 goat anti-rabbit/mouse, ThermoFisher Scientific). Following incubation, sections were rinsed 3x2 min in 0.1M PBS, mounted onto subbed glass slides, and were coverslipped using ProLong Antifade Reagent containing DAPI (ThermoFisher Scientific).

### Double Fluorescent *in situ* Hybridization

To obtain fresh frozen tissue for *in situ* hybridization, several Dlx1-CreER:Ai14 and Ascl1-CreER:Ai14 mice were selected following different time points of tamoxifen induction. Fresh tissue was harvested, rinsed in 0.9% saline, and submerged in Tissue-Tek O.C.T embedding media (VWR) in a cryomold container. This mold was quickly placed in 2-methyl butane (Sigma), which then was placed in liquid nitrogen until the OCT block was frozen solid. The resulting frozen tissue was stored in −80°C freezer before cryoosectioning.

Two-probe fluorescent *in situ* hybridization was performed on fresh frozen 12-µm thick cryosections of P28 mouse brains using the QuantiGene ViewRNA ISH assay from Affymetrix. Briefly, tissue was treated with formaldehyde fixation, protease digestion, target and label probe incubation, and buffer washing according to the QuantiGene Affymetrix protocol. Tissue was then co-labeled with fluorescent Nissl and imaged using a DeltaVision epifluorescent microscope (details below).

### Imaging and Serial Reconstruction of Whole Brains

Fate-mapped neurons were imaged from serially mounted sections on a Zeiss LSM 780 confocal microscope (CSHL St. Giles Advanced Microscopy Center) and Zeiss LSM 510 confocal microscope with ZEN software (Ragon Institute of MGH, MIT and Harvard), including all featured images with immunostaining. Volumetric whole brain imaging of native fluorescent signal was completed with a 5-µm Z-step using previously described methods for serial two-photon tomography (Kim et al., 2015, Ragan et al., 2012). Briefly brains were embedded in oxidized agarose and placed in a motorized stage in a TissueCye 1000 (Tissuevision). Whole brains were then imaged as 12 (X) x 16 (y) tiles with 1 µm x 3 1 µm resolution for 270 Z sections (50-µm interval) or 1500 Z sections (5-µm interval). Image datasets were stitched and reassembled by custom built software and viewed as stacks or 3D volumes. Image processing was completed using ImageJ/FIJI software with all alterations applied only to entire images. Three-dimensional image stacks of striatal cells from *in situ* data were acquired at room temperature with a DeltaVision restoration microscope system consisting of an Olympus IX71 inverted microscope equipped with Olympus 20X/0.75 U Apo 340 dry objectives and a Photometrics coolsnap HQ2 camera configured at 0.1122 µm/pixel. Filters used were 430/24 ex/470/24 em and 645/30 ex/705/72 em. Z-stack images of cells of cells were subjected to deconvolution by using Deltavision softworx 6.5.1 software (Applied Precision). The maximum projection or the middle optical section of the deconvolved sections was exported to TIFF format and then processed by FIJI/ImageJ for the final images.

### Quantification and Data Analysis

Manual scoring and counting of SPN cell types in striatal compartments was completed using Neurolucida 11 software either directly from slides on an Olympus Bx51 microscope or from serially collected Z-stack tile scan images (confocal or epifluorescent). Specifically, striosomes were identified positively as groups of clustered cells with thick spinous dendrites co-localizing to MOR1-positive zones and in negative by the absence of both Calbindin 28K (CB) and CDG1 co-localization, which are specifically expressed in the matrix compartment of the striatum. Unless otherwise indicated, each quantified sample consists of cell counts from 6 representative rostral-caudal coronal sections of striatum per hemisphere, with multiple (2–10) brains from at least two litters sampled in all cases. Cell densities in striosome and matrix compartments were calculated in two in areas bounded by drawing contours around the Mor1^+^ striosomes or CDG1^+^ matrix and counting the number of RFP^+^ cells within each compartment, averaging the cell density from replicate sections as done for all other cell counts. Comparisons of striosome content in rostral (Bregma 1.54–0.5 mm) versus caudal (Bregma 0.45–0.82 mm) sections through the striatum at E12.5 TM (Ascl1-CreER) and E13.5 TM (Dlx1-CreER) were quantified in counts in serial sections (on average 42 sections of 55-µm thickness). Rostral/caudal levels were defined according to the 4^th^ edition of the Mouse Brain in Stereotaxic Coordinates atlas (Paxinos and Franklin, 2001). Finally, to quantify the co-localization of BrdU with RFP^+^ cells, the ratio of BrdU^+^RFP^+^ cells in striosomes to total RFP^+^ cells in striosomes was determined for a minimum of three different brains of two different litters in all cases. Unpaired or paired Student’s t-tests were used to analyze data with two groups (see each figure legend for details), and one-way ANOVA and Tukey’s honest difference significance tests were used for unbiased comparisons of multiple time-points within a series. Results were displayed graphically and statistics completed using Graphpad Prism software.

## SUPPLEMENTAL FIGURE LEGENDS

**Figure S1.**
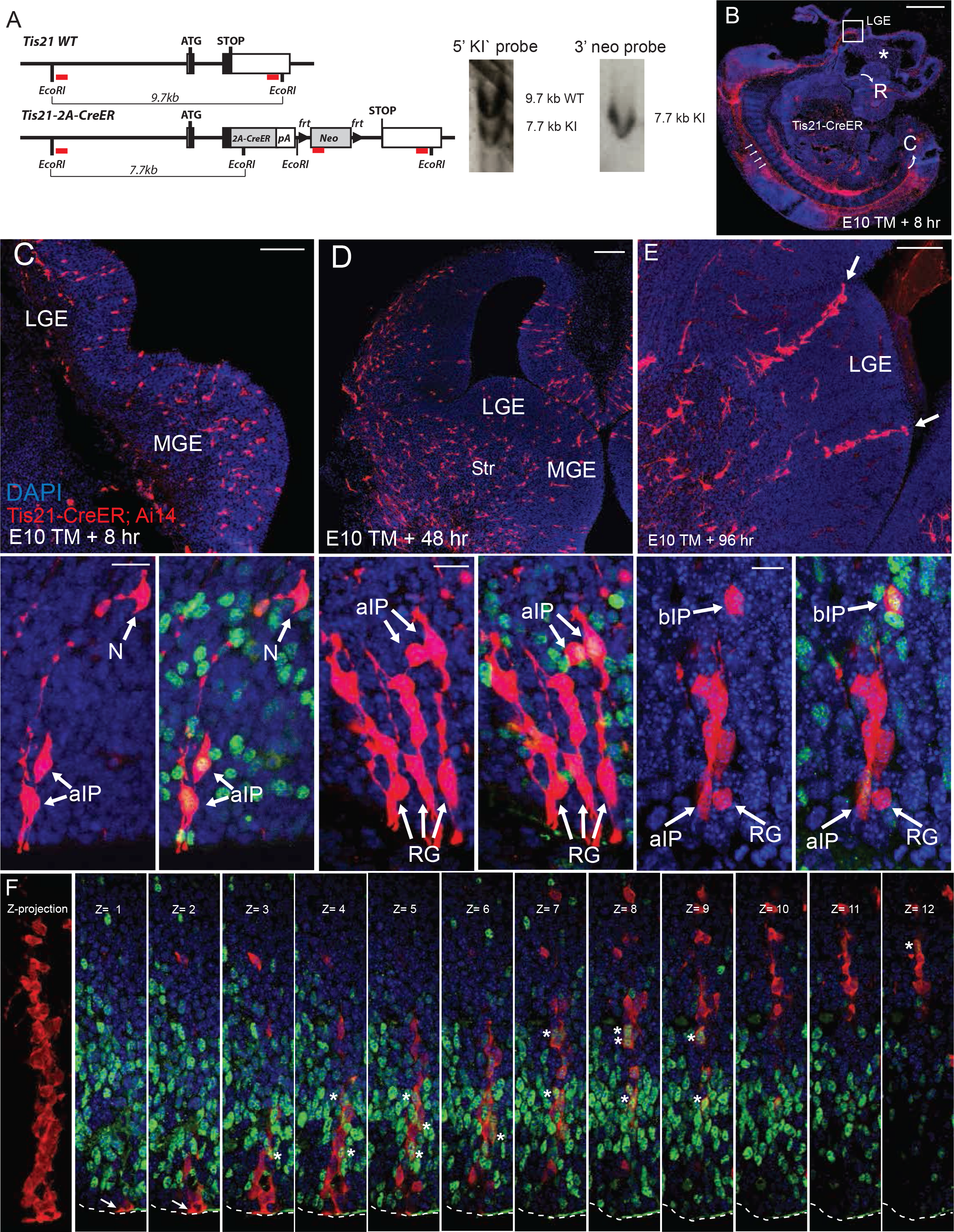
The Tis21-CreER Driver Allows Genetic Fate Mapping of Neurogenic RGs and IPs throughout LGE Neurogenesis, Related to Figure 1. (A) Gene targeting strategy and southern blot confirmation of a targeted *Tis21-2A-CreER* ES cell clone. A 2A-CreER-polyA cassette was inserted in-frame before the stop codon of the Tis21 gene. Frt-neo-frt cassette was removed by breeding with Flp transgenic mice. Red bar: southern probe fragment. (B) E10 TM Pulse-chase of 8 hours in *Tis21-2A-CreER;Ai14* embryos labels neurogenic progenitors throughout the neural tube, including all germinal domains from rostral (R) regions such as the telencephalon (*) down to the caudal (C) regions of spinal cord (arrows). (C-E) E10 TM pulse-chase in *Tis21-2A-CreER;Ai14* embryo labels a sparse assortment of RGs and aIPs in the LGE, visible at 8 hours (C). These progenitors subsequently produce neurons that migrate into the developing striatum, seen at 48 hours (D) and 96 hours (E) post-induction. Bottom panels depict examples of RGs, aIPs, bIPs and postmitotic neurons (N) during this period of LGE neurogenesis. (F) The left most panel shows the projected image of a particular large clone at 96 hours following E10.5 TM induction. Remaining panels show a montage of individual optical sections across this isolated cluster. The RG maintains end-feet at ventricle surface (arrow) and has generated multiple Ascl1^+^ bIP (asterisks) throughout its vertical axis along the expanded SVZ by E14.5, suggesting multiple rounds of RG division and significant proliferation of bIPs during the 96 hour period. Scale bars: 1 mm in A; 100 µm in top of C, D, E; 20 µm in bottom of C, D, E;

**Figure S2.**
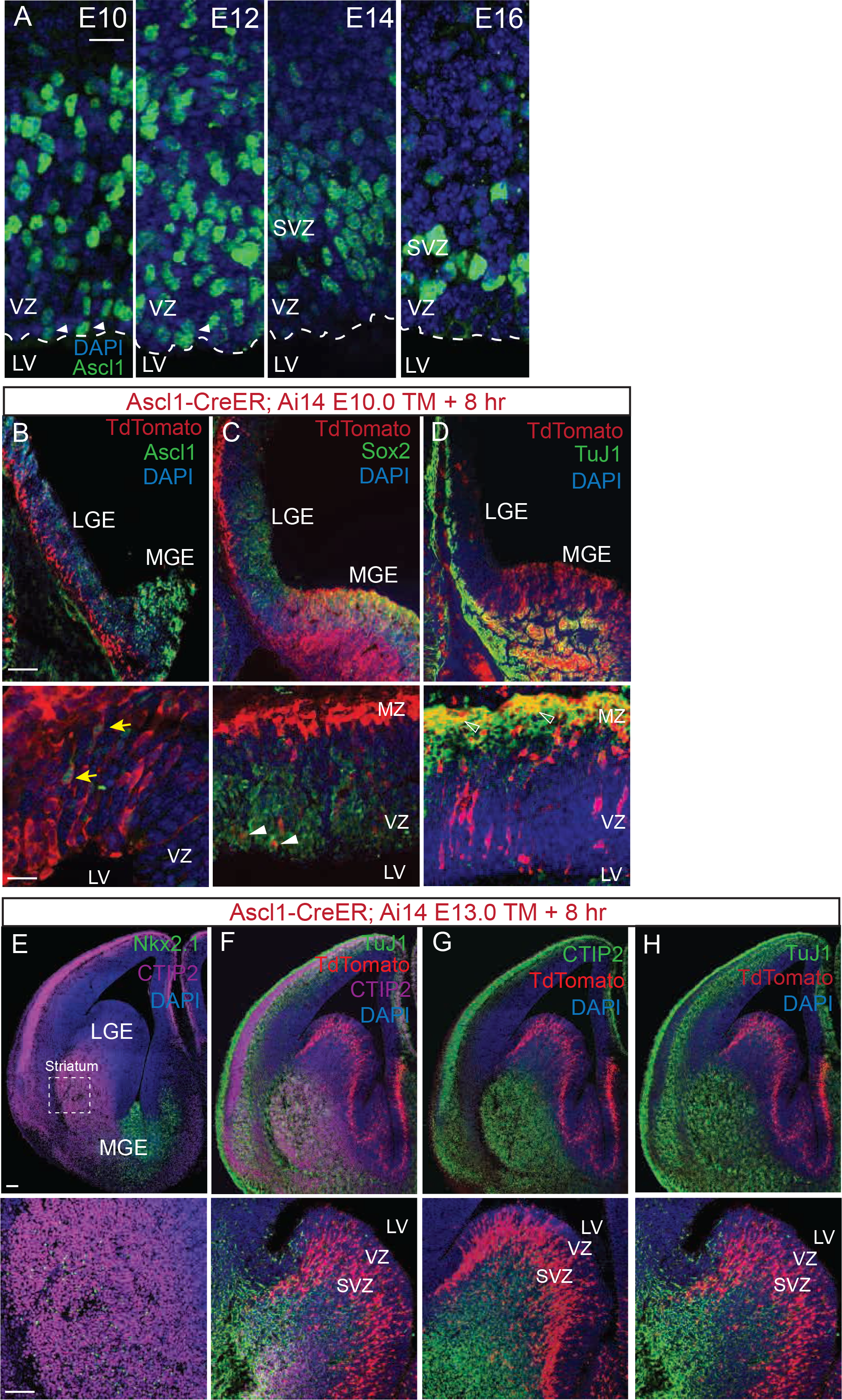
The *Ascl1-CreER* Driver Targets Early aIPs and Late bIPs in the LGE, Related to Figure 2. (A) High magnification images showing ASCL1 immunoreactivity (green) in LGE progenitors at E10, E12, E14, and E16. Note the progressive shift of Ascl1 expression from VZ to SVZ starting at E12 and restriction to SVZ by or before E14. Arrowheads indicate putative aIPs with somas at the ventricular surface at E10. Blue: DAPI. (B) Low (top) and high (bottom) magnification images showing ASCL1 immunoreactivity (green) in progenitors marked by *Ascl1-CreER;Ai14* E10 TM induction (red, yellow arrows) with 8 hour interval before harvesting. Blue: DAPI. (C) Low (top) and high (bottom) magnification images showing expression of the multipotency factor Sox2 (green) in progenitors marked by Ascl1-CreER;Ai14 E10 TM induction (red, arrowheads). Blue: DAPI; yellow: merge of red and green. (D) The marker of postmitotic migrating neurons (open arrowheads) BIII-tubulin 1 (TuJ1, green) does no co-localize with aIPs marked by *Ascl1-CreER;Ai14* E10 TM induction (red). Blue: DAPI; yellow: merge of red and green. (E) The MGE-specifying homeodomain factor Nkx2.1 (green) and neuronal marker CTIP2 (purple) are expressed respectively in non-overlapping populations of local GABAergic interneurons and spiny projection neurons in striatum at E13.5. Blue: DAPI. (F) bIPs and aIPs marked by *Ascl1-CreER;Ai14* E13 TM induction remain in the SVZ at 8 hours following TM induction, largely separated from the previously born postmitotic neurons expressing CTIP2 (purple) and BIII-tubulin1 (TuJ1, green) Blue: DAPI. (G) Newly postmitotic SPNs deriving from progenitors marked by *Ascl1-CreER;Ai14* E13 TM acquire expression of CTIP2 (green) upon reaching the striatum. Blue: DAPI. (H) Postmitotic SPNs deriving from progenitors marked by *Ascl1-CreER;Ai14* E13 TM acquire expression of TuJ1(green) upon reaching the striatum. Blue: DAPI. Scale bars: 20 µm in A; 100 µm (top), 10 µm (bottom) in B; 100 µm in E.

**Figure S3.**
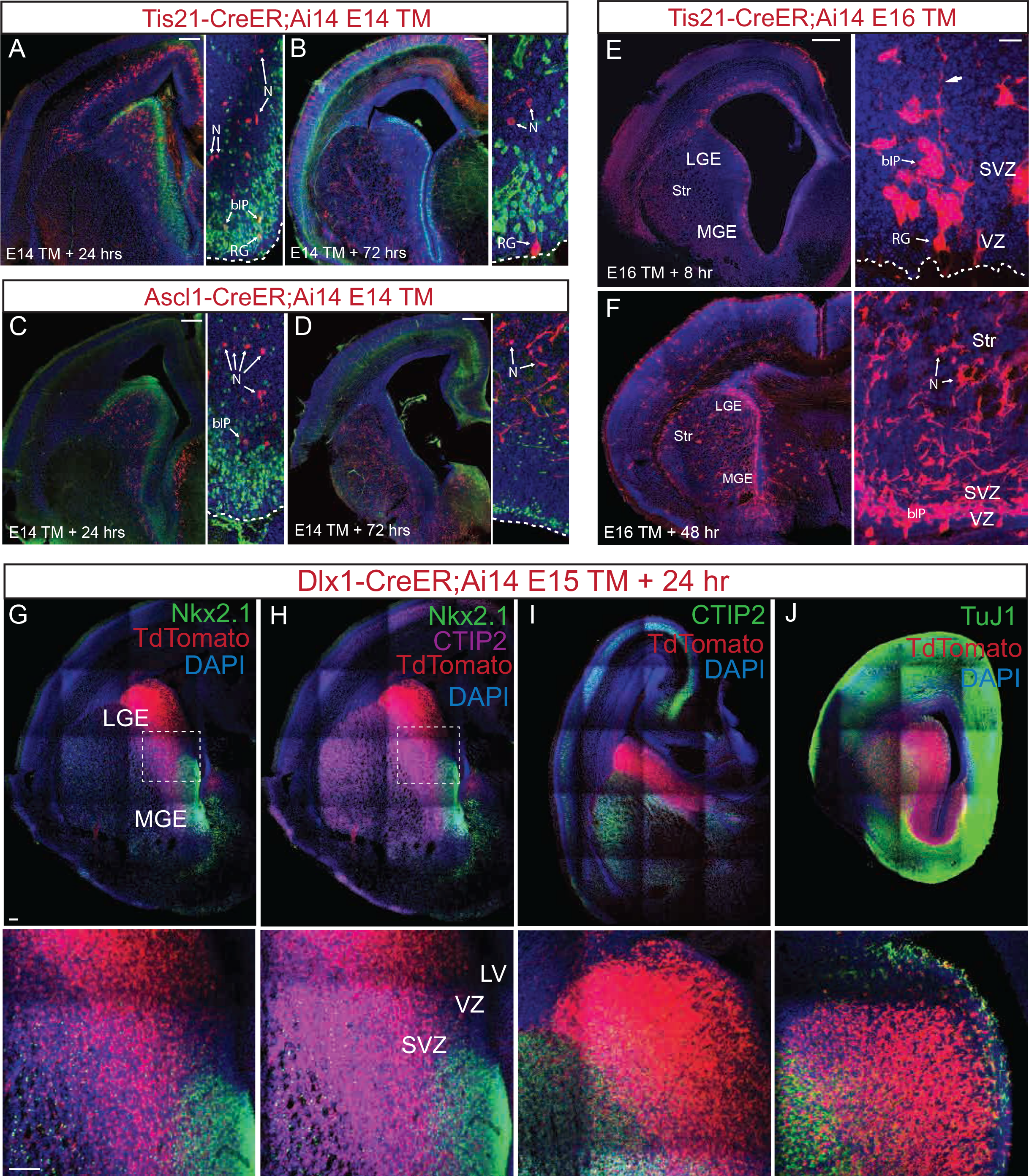
Characterization of Late LGE Progenitors, Related to Figure 3. (A and B) Pulse-chase analysis of *Tis21-CreER;Ai14* embryo at 24 (A) and 72 (B) hours following TM induction at E14.5. Self-renewing RGs remained at VZ and continued to generate bIPs and neurons up to 72 hours later. (C-D) Pulse-chase analysis of *Ascl1-CreER;Ai14* embryo at 24 (C) and 72 (D) hours following TM induction at E14. Labeled bIPs were deleted in VZ and SVZ by 24 hours. Differentiating SPNs in MZ were abundant at 72 hours. (E and F) Eight-hour pulse-chase in E16 *Tis21-CreER;Ai14* embryo continued to label neurogenic progenitors in LGE and throughout the ventricle wall (E). Postmitotic SPNs derived from these progenitors and migrating into the developing striatum were abundant 48 hours (F, arrows). At higher magnification, RG, bIP, and postmitotic neurons can be identified by their location and morphology in VZ, SVZ, and MZ. (G) TM induction at E15 in *Dlx1-CreER;Ai14* embryos labeled dense populations of LGE IPs and subset MGE progenitors, resolved by co-localization with anti-Nkx2.1 (green). (H) LGE- and MGE-derived neurons marked by E15 TM induction in *Dlx1-CreER;Ai14* mice intermixed in striatum as soon as 24 hours following induction. (I) Postmitotic SPNs derived from E15.5 Dlx1^+^ bIPs acquire expression of CTIP2 (green) upon reaching the striatum. Blue: DAPI. (J) Postmitotic SPNs derived from E15.5 Dlx1^+^ bIPs acquire expression of TuJ1 (green) upon reaching the striatum. Blue: DAPI. Scale bars: 300 µm in A-D, E (left); 20 µm in E (right); 100 µm in G.

**Figure S4.**
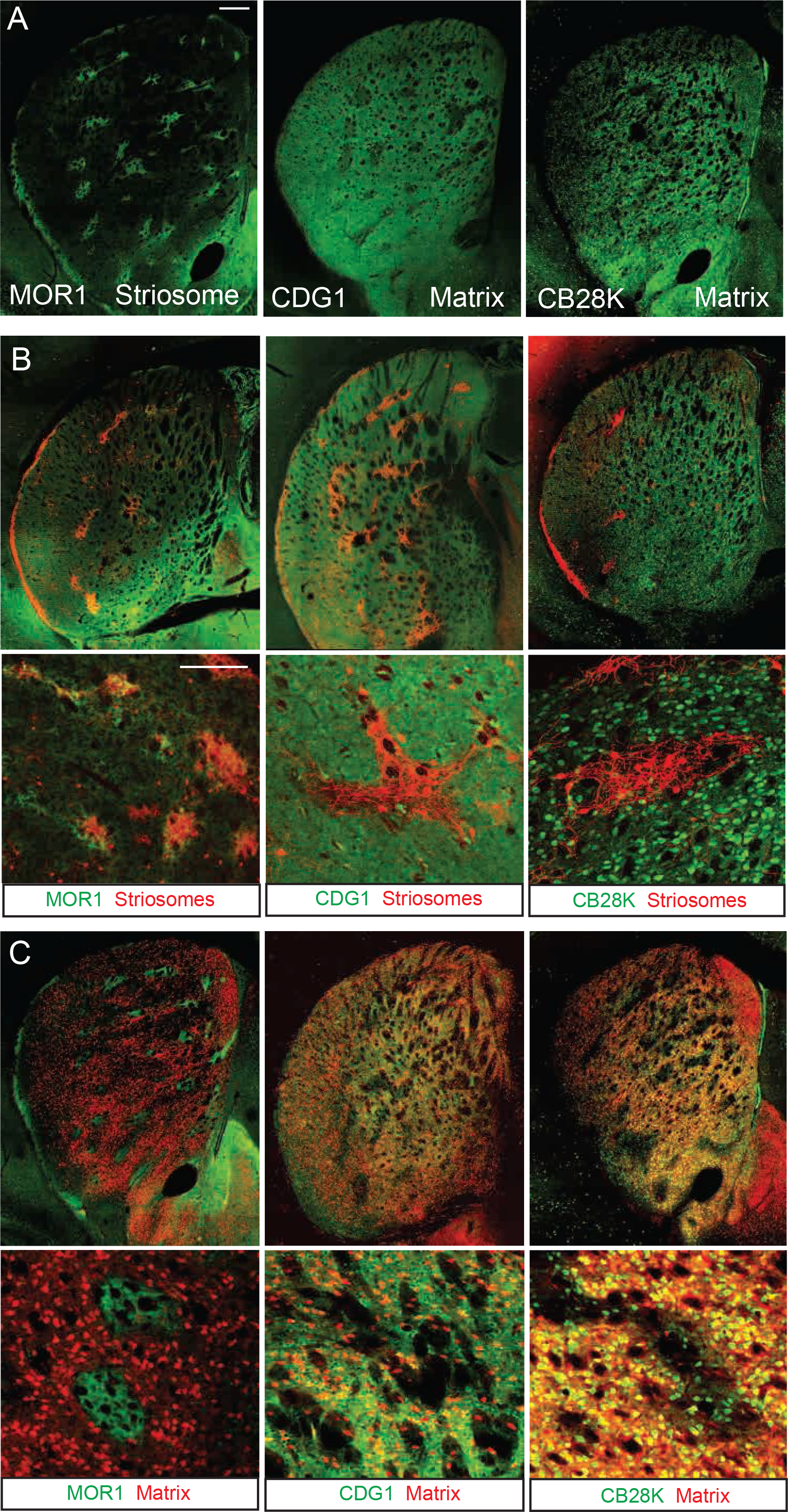
Co-Labeling of Fate-Mapped SPNs with Striatal Compartment Markers, Related to Figures 2, 3, and 5. (A) Examples of MOR1 immunostaining (left) as the striosomal marker and CDGI (middle) and Calbindin 28K (right) immunostaining as the matrix markers. (B) Co-labeling of striosomes (red) following E10.5 TM induction in *Ascl1-CreER;Ai14* mice with, from left to right, anti-MOR1, anti-CDGI, and anti-CB28K immunostaining, respectively (green). (C) Co-labeling of matrix following E17.5 TM induction in *Dlx1-CreER;Ai14* mice with, from left to right, anti-MOR1, anti-CDGI, and anti-CB28K immunostaining, respectively (green). Scale bars: 300 µm.

**Figure S5.**
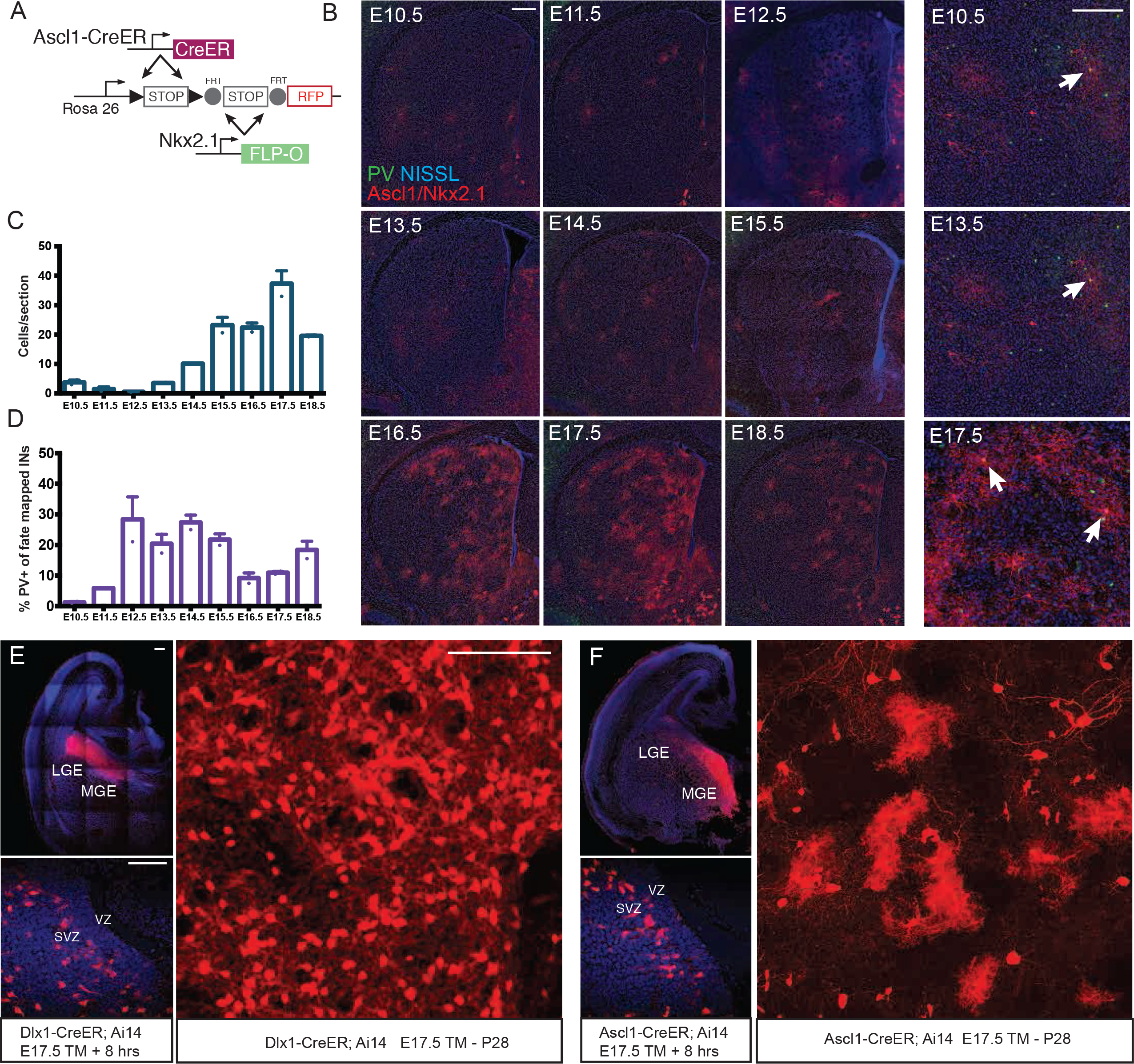
Striatal GABAergic Interneurons Arise from Progenitors Distinct from Those That Give Rise to SPNs, Related to Figure 5. (A) Inducible intersectional genetic fate-mapping strategy for fate mapping the striatal interneurons deriving from MGE and preoptic area based on the targeting of progenitors that express both Ascl1 and Nkx2.1, either simultaneously or sequentially. (B) Representative striatum panels of TM inductions in *Ascl1-CreER;Nkx2.1-ires-Flp-O;Ai65* mice, showing a sparse assortment of interneurons in cohorts captured by TM inductions from E10.5 to E15.5 (red) and denser populations from E16.5 to E18.5. Green: parvalbumin. blue: Nissl. Right column: parvalbumin-positive cells (arrows) at E10.5, E13.5, and E17.5. Scale bars: 300 µm (left), 50 µm (right). (C) Histogram of striatal interneuron production from intersectional fate mapping of MGE progenitors from E10.5 to E18.5. Eighty-five percent of all interneurons derive from progenitors captured by TM inductions E15-E18, whereas only 5% (aggregate) come from TM inductions E10-E12. Error bars, SEM. (D) Proportional histogram of parvalbumin immunoreactive interneuron content in intersectional fate mapping of MGE progenitors from E10.5 to E18.5. (E) Dlx1^+^ SVZ progenitors are arranged in a dorsally focused gradient in the E17 LGE, and give rise to a dense population of matrix SPNs. (F) Ascl1^+^ SVZ progenitors are arranged in an opposing ventrally focused gradient to that of Dlx1^+^ progenitors in the E17 LGE, giving rise to glia cells and presumably MGE-derived local striatal interneurons in preference to SPNs. Scale bars: 100 µm in E, F.

**Figure S6.**
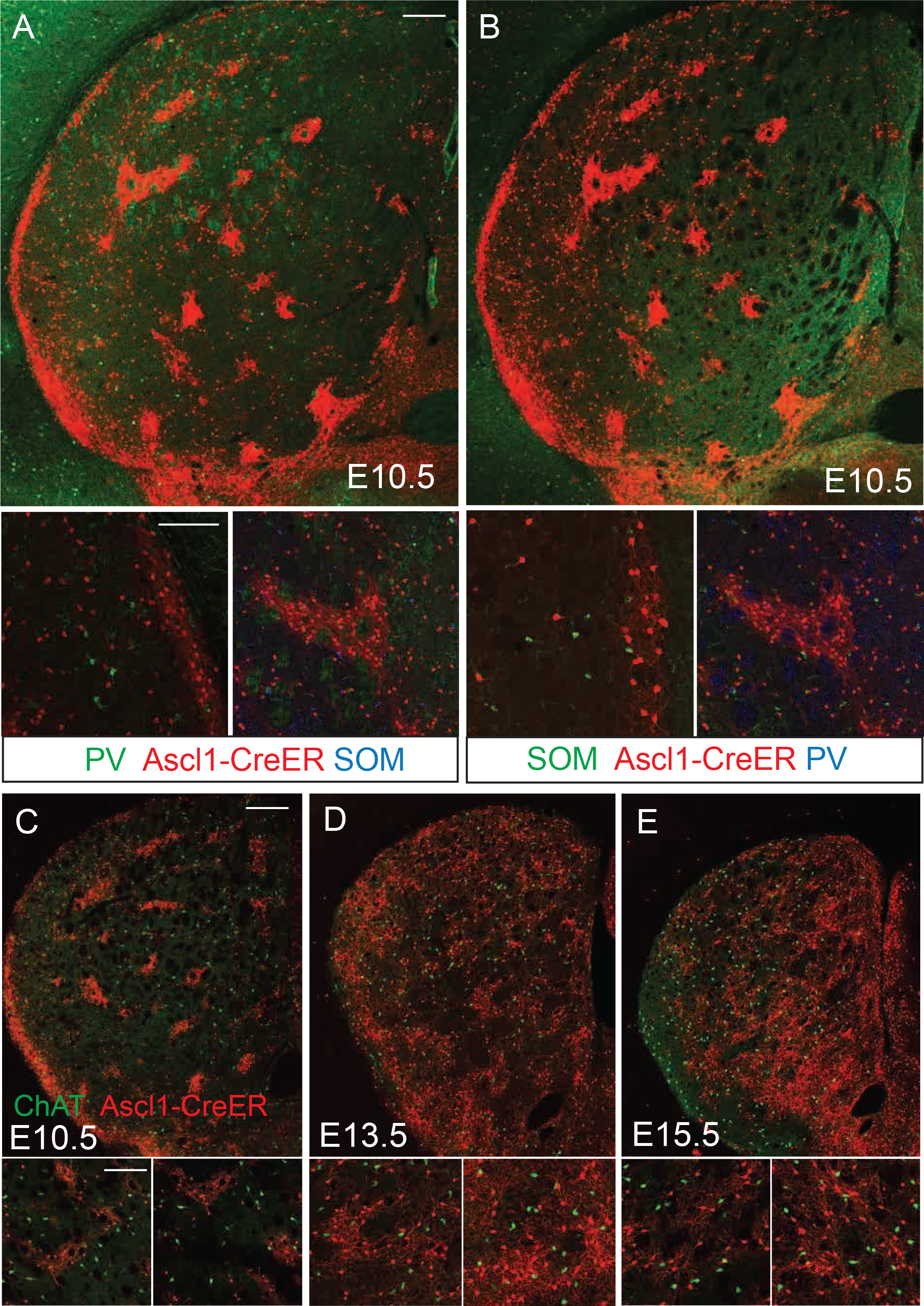
Cholinergic Interneuron Content in Striosome Fate Mapping, Related to Figure 5. (A) Immunolabeling of parvalbumin (PV, green) and somatostatin (SOM, blue in subpanel) does not overlap with fate-mapped striosomes following E10.5 TM induction in *Ascl1-CreER* mice. (B) Immunolabeling of somatostatin (SOM, green) and parvalbumin (PV, blue in subpanel) does not overlap with fate-mapped striosomes following E10.5 TM induction in *Ascl1-CreER* mice. (C-E) Immunolabeling of choline acetyltransferase (ChAT) expression in cholinergic interneurons (ChAT, green) does not overlap with fate-mapped striosome or matrix deriving from the *Ascl1* lineage following TM inductions at E10.5 (C), E13.5 (D), or E15.5 (E) in *Ascl1-CreER* mice (*Ascl1-CreER;Ai14*, red). Scale bars: 100 µm.

**Figure S7.**
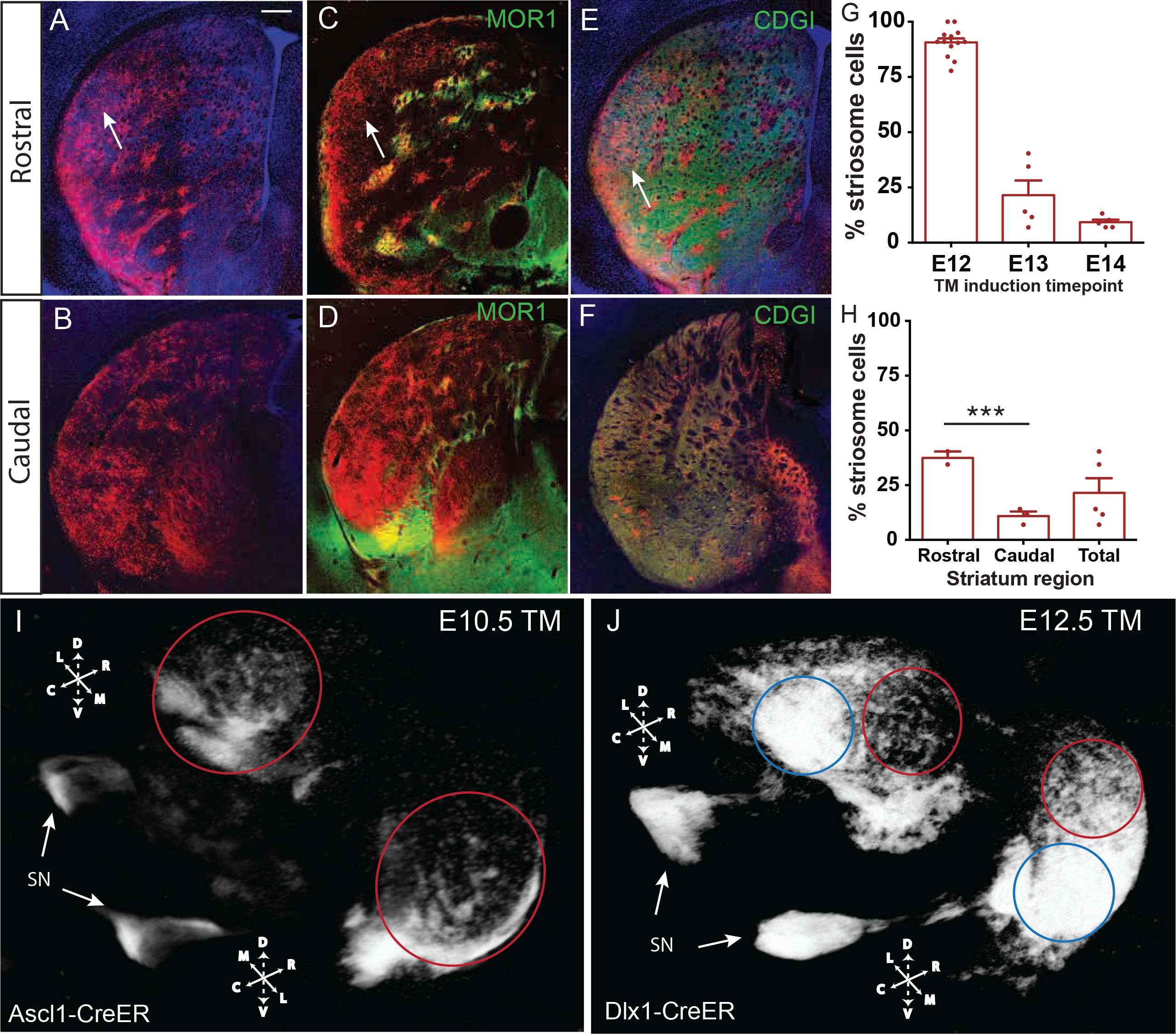
Temporal Order of Neurogenesis in LGE Translates into a Spatial Sequence of SPN Settlement in Striatum, Related to Figure 5 and 6. (A and B) Distribution of fate-mapped SPNs in rostral (A) and caudal (B) striatum following E12.5 TM induction in *Ascl1-CreER;Ai14* mice. Arrow: lateral infiltration of the earliest matrix SPNs to populate rostral striatum. Scale bar: 300 µm. (C and D) Co-labeling of striosomes in rostral (C) and caudal (D) striatum by anti-MOR1. (E and F) Co-labeling of matrix in rostral (E) and caudal (F) striatum by anti-CDGI. (G) Decline in S cell production from E12.5 to E14.5 in *Ascl1-CreER;Ai14* mice. Error bars, SEM. (H) Pooled quantification of S cell content in rostral versus caudal striatum following E12.5 TM induction in *Ascl1-CreER;Ai14* mice or E13.5 TM induction in *Dlx1-CreER;Ai14* mice. ***p < 0.0001. (I) Serial two-photon tomography of a P28 brain following E10.5 TM induction in a *Ascl1-CreER;Ai14* mouse revealed a dense shell of S cells encapsulating the lateral border of striatum and sparsely distributed striosomes throughout both the caudal and rostral extent of the striatum (red circle), with axon projection to the SNc (arrows). Dorsal-ventral (V-D), lateral-medial (L-M) and rostral-caudal (R-C) axes are indicated. (J) Serial two-photon tomography of a P28 brain following E13.5 TM in a *Dlx1-CreER;Ai14* mouse shows a sharp transition from matrix (blue circle) to striosomes (red circle) in caudal compared to rostral regions of striatum. The birth date-dependent lateral to medial settling occurred in two staggered waves 9 visualized in both Ascl1 and Dlxl1drivers - the first for S cells between E10-E13 and the second more prominent gradient for M cells between E13-E17 (see Figure 5).

